# Design of proteins by parallel tempering in the sequence space

**DOI:** 10.1101/2025.03.13.643021

**Authors:** Preet Kalani, Vojtěch Spiwok

**Affiliations:** Department of Biochemistry and Microbiology, University of Chemistry and Technology, Prague, Technická 3, Prague 6, 166 28, Czech Republic

**Keywords:** protein design, ESMfold, machine learning, Monte Carlo, parallel tempering, replica exchange

## Abstract

Design of new proteins is often formulated as an optimization task. An amino acid sequence is characterized by an energy, and this energy is sampled and minimized. Here, we use a parallel tempering algorithm to accelerate this task. A series of 100- or 200-residue proteins was designed using a modified Evolutionary Scale Modeling design module to maximize the confidence in structure prediction and globularity and minimize the surface hydrophobic residues. We show that parallel tempering is a viable alternative to Monte Carlo sampling and simulated annealing or related energy-based protein design methods, especially in the situation where a continuous flow of designed sequences is desired.

## 1 INTRODUCTION

The design of new proteins has become a viable strategy to obtain new functional (e.g. binding or catalytic) proteins, as an alternative to the exploration of natural sources and protein engineering ^1^. Many of protein design approaches are based on iterative modification of the amino acid sequence from, for example, a random sequence, until desired properties of the protein are reached. Such protein design campaigns are based on two components.

First, we need a function to assess an amino acid sequence to determine whether it folds into the desired 3D structure and fulfills other requirements for the target application. These functions may be energy-based (based on potential or free energy) versus knowledge-based (based on known sequences and 3D structures), traditional physics versus machine learning, or free (favoring any compactly folded 3D structure) versus targeted (favoring only proteins with desired 3D structure, symmetry, etc.).

Second, it is necessary to employ an algorithm that samples various sequences and optimizes the above described function. Again, various approaches can be used in this stage, including physics-inspired algorithms such as the Monte Carlo method and simulated annealing ^2–5^ as well as machine learning methods ^6,7^.

Despite recent success in this field, the success rate of protein design projects is not 100 %. Usually, multiple proteins must be designed to obtain one that passes the experimental evaluation. Furthermore, some features can be easily controlled during the design process, whereas some other properties are very difficult to control, thus requiring intensive experimental testing. Current protein design methods usually provide one designed sequence in one optimization run, rather than a continuous flow of designed sequences. Methods providing a continuous flow of designed sequences may be more suitable in the situation when multiple protein design candidates must be tested.

Monte Carlo ^8^ and Monte Carlo simulated annealing have been widely used to evolve sequences in protein design ^2–5^. In general, the sequence undergoes small changes that can be accepted or rejected. A change leading to a protein with better properties (lower energy *E*) is always accepted. A change leading to a protein with worse properties is either accepted or rejected with the probability *p* calculated by Metropolis criterion ^8^:

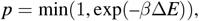

where *E* is the energy and *β* is an inverse temperature (1/*kT*, where *k* is Boltzmann constant and *T* is temperature). The fact that even sequences with higher energy can be accepted ensures that the system can escape a local minimum and proceed towards the global one (the optimal sequence in the case of protein design).

High temperature accelerates sampling, whereas low temperature favors low-energy states. Simulated annealing uses a predefined change of temperature to ensure that a wide sequence space is sampled at high temperature and, at the same time, a sequence close to the minimum of the sampled space can be identified at low temperatures. Most applications of Monte Carlo simulated annealing in protein design use a schedule with a single monotonous or stepwise decrease of temperature. Unfortunately, one such run leads to one designed protein, which may be inefficient in the situation when multiple designed proteins must be tested.

Here, we explore the application of the parallel tempering algorithm ^9^ (also known as the temperature replica exchange algorithm) in protein design.

Parallel tempering is a popular method used in modeling physical and chemical systems. It can be combined with Monte Carlo method or molecular dynamics simulation. In general, the system studied is simulated by one of these methods in multiple replicas that differ in temperature. Occasionally, an attempt is made to exchange systems simulated at different temperatures. The probability of the exchange is calculated as:

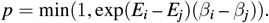

The indexes *i* and *j* refer to the temperature IDs. Next, a uniformly distributed random number between 0 and 1 is drawn. If this number is lower than *p*, replicas are exchanged, i.e. simulation of *i*-th system continues at *j*-th temperature and vice versa. Such replica exchange attempts are made periodically.

The application of parallel tempering to protein design is schematically depicted in Figure 1. In the beginning, a series of random protein sequences is generated (the first column in Figure 1). Next, they are subjected to single-temperature Monte Carlo sampling, each at a different pre-selected temperature. After a predefined number of sampling steps (gray arrows in Figure 1), a replica exchange attempt is made.

**FIGURE 1.**
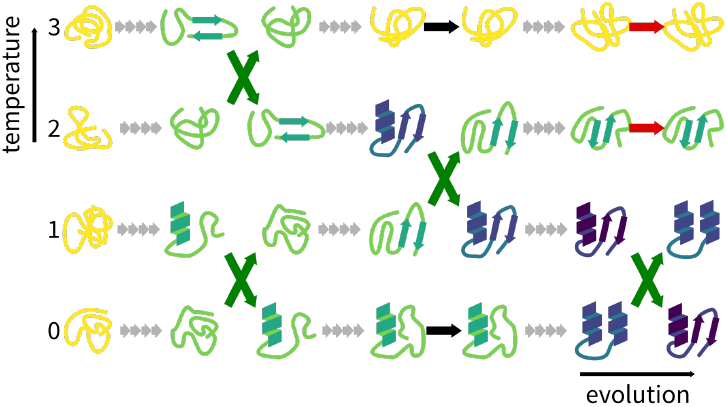
Schematic depiction of protein design by parallel tempering.

A sequence corresponding to an unstable protein (with high energy *E*) is likely to be exchanged for higher-temperature replicas and thus climb on the temperature scale. At high temperature, it can (due to faster sampling) reach a sequence corresponding to a stable protein (with low energy *E*). This sequence tends to decline on the temperature scale. Stable sequences, therefore, accumulate at low temperature. Parallel tempering is in general significantly more efficient in exploration of different states of the system and in optimization than a series of equally long single-temperature sampling or simulated annealing runs.

Here, we combine parallel tempering with protein design. Protein sequences were optimized to provide stable folded proteins. Parallel tempering was used as part of this algorithm. Our approach is based on Evolutionary Scale Modeling (ESM) ^10^, which is a large language model used in the 3D structure prediction tool ESMfold ^11^. The modified version of the “protein programming language” of ESMfold ^3^ was used as an engine that generates and scores sequence candidates and runs the Monte Carlo method. The “protein programming language” makes it possible to easily define properties of the desired protein (e.g. compactness, few hydrophobic residues on the surface, secondary structure content, or similarity to a reference 3D structure).

We demonstrate our protocol in the free (untargeted) design of proteins with 100 and 200 amino acid residues. We show that our method is efficient in the continuous design of folds differing in overall shape and secondary structure composition.

## 2 RESULTS

The design of proteins by parallel tempering used a modified version of ESMfold, namely its protein design module (“protein programming language”) ^3^. This module uses the Monte Carlo simulated annealing method to minimize a score calculated from the sequence. This score can be defined by a modular language to combine features of the designed protein. Using a Python program, we introduced parallel tempering into this code. The initial sequences were generated randomly with the same probability of all amino acids except cysteine. They were subjected to Monte Carlo sampling with replica exchanges made, in most cases, every 100 Monte Carlo steps.

Figure 2 presents the progress in the design of 100-residue proteins. Figure 2A shows the profile of replica exchanges. Each color represents one replica. The selected replica (starting from temperature ID 16) is highlighted by a thick line. It is clear that replica exchanges were frequent and there were no problematically selected temperatures. Figure 2B shows the evolution of the score. At first, the amino acid sequences were set randomly with equal amino acid preferences. Therefore, the score was very high at all temperatures, indicating the unstructured nature of the proteins. After relatively few steps (approx. 5,000 steps, 50 replica exchange attempts), stable designs were formed at low temperatures. The proteins remained poorly structured at high temperatures. Figure 2C shows evolution of mean C*α* predicted local distance difference test (pLDDT) values. It shows a trend similar to Figure 2B. This figure also highlights the demultiplexed design profile (red trace) with representative 3D structures predicted by ESMfold.

**FIGURE 2.**
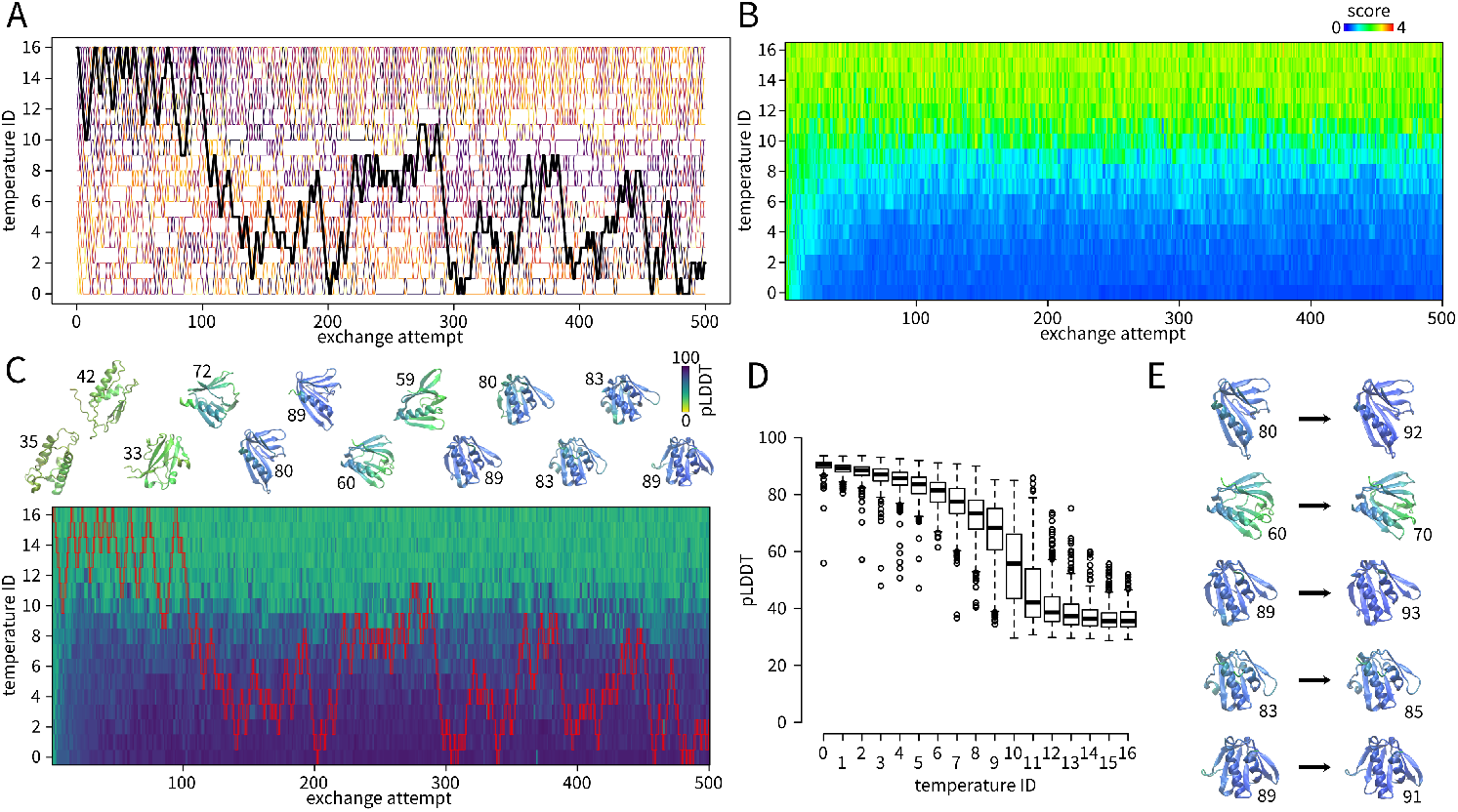
Design of 100-residue proteins by parallel tempering. A – replica exchanges. Selected replica starting from temperature ID 16 is highlighted in tick. B – evolution of the score. C – evolution of mean C*α*-pLDDT. The replica starting from temperature ID 16 is highlighted by red boxes. Snapshots of ESMfold models from demultiplexed trace are depicted at top (colored by pLDDT, numbers indicate the mean C*α*-pLDDT). D – boxplot of the mean C*α*-pLDDT at different temperatures (first 50 replica exchange attempts skipped, sampled at replica exchange points). E – refinement of models highlighted in C by simulated annealing (numbers indicate the mean C*α*-pLDDT).

Demultiplexing (also referred to as demuxing) is a reconstruction of a continuous sequence evolution regardless of the temperature. For example, the red profile Figure 2C started at the temperature number 16. In the next replica exchange attempt (after 100 Monte Carlo steps), it was exchanged for the replica at temperature number 15, and so forth. The demultiplexed evolution thus joins structures at temperatures 16 in step 0, 15 in step 100, etc.

Representative structures depicted in Figure 2C show that the initial protein with a random sequence was unstable (yellow on the pLDDT scale). After approximately 150 steps, a stable design (blue/purple in the pLDDT scale) was obtained. This was associated with a decrease in temperature. The resulting designs contained both *α*-helices and *β*-sheets. At replica exchange attempt approximately 280 it destabilized again (this time it was associated with a slight increase of temperature), and at replica exchange attempt approx. 300 it formed stable helix-rich designs. They remained stable with miner changes until the end of the run. The plots for the 17 replicas can be obtained as Supporting Information (Figures S1-S17).

Figure 2D shows a boxplot with mean C*α*-pLDDT values. It further supports the notion that designs at low temperatures are stable (pLDDT around 90 %), whereas they are unstable at high temperatures.

It is possible that the lowest temperature used in our replica exchange scheme is not low enough and that the designed proteins can be further optimized by reducing the temperature. To address this, we took snapshots in Figure 2C and subjected them to Monte Carlo simulated annealing optimization to reduce temperature. This refinement led to some improvement in the mean C*α*-pLDDT, between 2 and 12 percent points (Figure 2D).

The common problem of application of replica exchange methods is the low replica exchange probability. For this reason, we tested a higher number of replica exchange attempts. Replica exchange attempts were made every 20 steps, instead of every 100 steps. The results are presented in Supporting Information (Figure S18). The increase in replica exchange frequency did not significantly increase overall design efficiency.

After successful application to 100-residue proteins, we applied our pipeline to 200-residue proteins. The results are depicted in Figure 3. It is clear that our approach is efficient for larger proteins. It was necessary to use a larger number of replicas (24 instead of 17). Figure 3A shows that the replica exchange rate for temperatures 0–3 and 14–18 was lower than for 100-residue proteins, nevertheless it was still sufficient.

**FIGURE 3.**
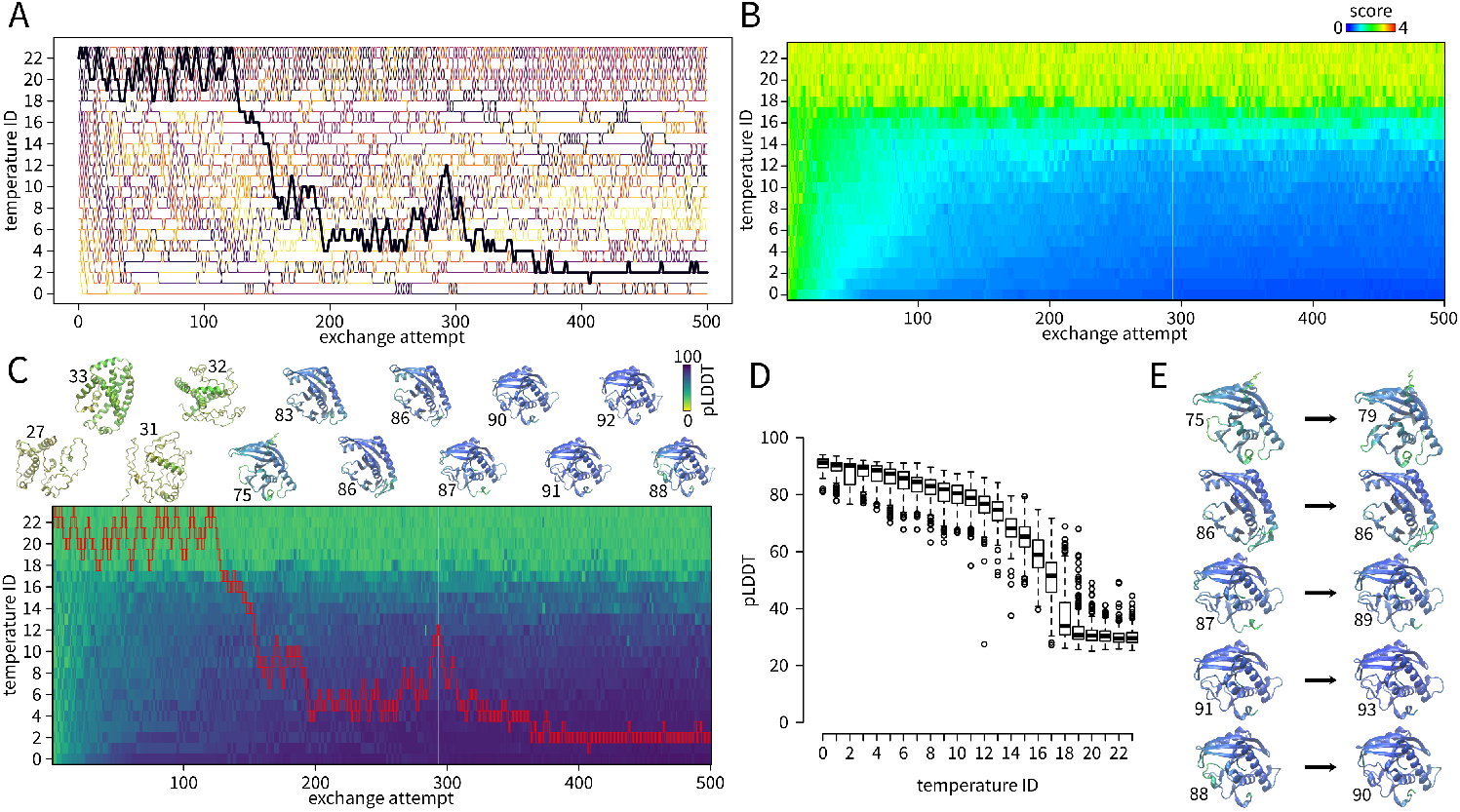
Design of 200-residue proteins by parallel tempering. A – replica exchanges. Selected replica starting from temperature ID 22 is highlighted in tick. B – evolution of the score. C – evolution of pLDDT. The replica starting from temperature ID 20 is highlighted by red boxes. Snapshots of ESMfold models from demultiplexed trace are depicted at top (colored by pLDDT, numbers indicate the mean C*α*-pLDDT). D – boxplot of the mean C*α*-pLDDT at different temperatures (first 50 replica exchange attempts skipped). E – refinement of models highlighted in C by simulated annealing (numbers indicate the mean C*α*-pLDDT).

As expected, a higher number of steps were needed for equilibration, i.e. to get to the state at which the distributions of energy at all temperatures become stable. Similarly to 100-residue proteins, the protein sequences sampled at the lowest temperature had favorable pLDDT values from ESM- fold (Figure 3C and D). Refinement of sequences by simulated annealing slightly increased pLDDT (by 0 – 4 percent points). Demultiplexed plots for all replicas can be obtained in Supporting Information (Figures S19-S42).

In order to further verify designed sequences, we predicted their structures using Alphafold 2^12^. Sequences of 100-residue proteins sampled at the lowest temperature (temperature ID 0, between replica exchange attempt 50 and the end, sampled every 10 replica exchanges) were submitted to Alphafold 2^12^ (the version for monomers) and the resulting pLDDTs were compared. The results are depicted in Figure 4. As expected, the Alphafold 2 pLDDT values were, in general, lower than the ESMfold pLDDT values because we optimized the ESMfold pLDDT. Alphafold 2 pLDDT values ranged from 36 to 91 % with a median of 62 %. In total 9 % had pLDDT greater than 80 and 22 % had pLDDT greater than 70, indicating a stable fold. The best Alphafold 2 score is highlighted in Figure 4.

**FIGURE 4.**
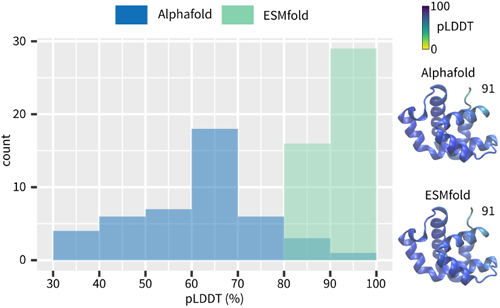
Histograms of pLDDT values of 200-residue proteins by Alphafold and ESMfold. The sequence with the highest Alphafold pLDDT score is depicted as Alphafold and ESMfold model colored by pLDDT (pLDDT indicated).

## 3 DISCUSSION

An interesting feature of our approach is that a demultiplexed replica presents an interesting evolution of the sequence and structure (Figure 2C and Figure 3C). Figure 2C nicely shows episodes of increased temperature associated with destabilization and decrease in temperature associated with the stabilization of designed proteins.

It must be kept in mind that in parallel tempering not only temperature influences protein stability, but also stability (energy *E*) influences temperature. High temperature naturally leads to faster sampling and destabilization. In contrast, low temperature favors low-energy states. In parallel, increasing *E* causes the replica to climb the temperature scale. Decrease of *E* leads to descent on the temperature scale. These two effects take place simultaneously.

The typical pattern of application of parallel tempering is a permanent increase and decrease in temperature. This is nicely visible in Figure 2. The protein is destabilized and stabilized again. This can be prolonged as necessary. Therefore, design by parallel tempering provides a continuous flux of designed sequences.

Many replicas in our protein design campaigns followed this patern; however, there were some demultiplexed replicas that stayed most of the time at low or high temperatures. Videos showing the evolution in Figure 2C and 3C can be found on Youtube (https://youtu.be/MOWU3L41lpM).

Parallel tempering is in general significantly more efficient than a series of equally long single-temperature Monte Carlo or molecular dynamics simulations. The gain in efficiency depends strongly on the studied system and the setup of the method. Yamamoto and Kob ^13^ used parallel tempering (replica-exchange) Monte Carlo method to accelerate the molecular structure of supercooled water. They observed that, depending on the setup, the parallel tempering is 10 to 100 times more efficient than a series of equally long simulations.

A critical step of the parallel tempering method is the choice of temperatures. They must be chosen in such a way that the highest temperatures allow for efficient sampling and, simultaneously, the lowest temperatures sample stable proteins. Neighboring temperatures *T*_*i*_ and *t*_*i*+1_ must be close enough so that the distributions of the energy values sampled *E*_*i*_ and *E*_*i*+1_ overlap. The overlap of the energy distributions ensures that *E*_*i*+1_ is occasionally lower than *E*_*i*_ or at least comparable, allowing for replica exchanges.

We used a trial-and-error approach to choose temperatures. In most cases, it is useful to use the constant ratio *T*_*i*+1_/*T*_*i*_. We used the highest temperature equal to 1 (in *kT* units), which we know from simulated annealing as a temperature at which sequences are sampled rapidly and leads to unstable proteins (data not shown). We set the ratio *T*_*i*+1_/*T*_*i*_ equal to 2 (temperatures 1, 0.5, 0.25, etc.) as a first attempt. It was necessary to reduce this factor for some temperature ranges to increase replica exchange rate.

We used different temperatures and different number of temperatures for the design of 100- and 200-residue proteins. This is analogous to the number of atoms in a system in a parallel tempering molecular dynamics simulation. Larger systems require a higher number of replicas because potential energy in a large system is averaged, and its variance is therefore lower. Lower variance means lower energy overlap of neighboring replicas, and thus lower replica exchange rate.

In addition to the size of the protein, the choice of optimized variables (pLDDT, pTM, surface hydrophobicity, etc.) and their weights may influence the choice of temperatures. However, temperatures can be set using a simple trial-and-error approach. In the future, we may develop a robust method for the choice of temperatures.

Parallel tempering has been extremely successful in connection with molecular dynamics simulation to model the folding of various small fast-folding proteins ^14^. A fundamental difference between parallel tempering in protein design and protein folding simulation (or any other biomolecular simulation) is the fact that the former aims at finding the global or close to the global minimum of *E*, whereas the latter aims at determining the distribution of states at the biologically relevant temperature. In protein design, we want to identify the sequence with the lowest possible *E*. In contrast, in protein folding simulations, we want to determine the fraction of states (e.g. unfolded and folded) at the biological temperature.

The choice of the lowest temperature in parallel tempering simulation of protein folding is usually straightforward and it is the biological temperature. In our application, we tried to overcome the fact that we cannot set the lowest temperature to 0 K by applying simulated annealing to selected designs (Figure 1E and 2E). The temperature dropped in these simulated annealing runs below 10^−6^*kT*. In general, there was an improvement in pLDDT.

The advantage of parallel tempering is its inherent parallelism. The design in each replica can proceed independently on the others, so they can be performed at different nodes of a parallel computer. Communication of nodes is necessary only in replica exchange attempts. Because we did not have access to a parallel computer with 17 or 24 GPUs, we performed all simulations in a serial manner. Users of parallel clusters may use a combination of message passing interface (MPI) with Python to parallelize the task.

The proteins designed in this article were not verified experimentally, i.e. by recombinant production of designed proteins and their structural characterization. We understand that this may discourage potential users of our approach. However, we believe that the quality of protein design campaigns is mainly determined by the quality of scoring of the designed sequences, rather than by the optimization method.

To our knowledge, the protein design engine used in this work (“protein programming language”) ^3^ has not been verified experimentally. A similar engine, also based on ESMfold, (“language model design”) ^2^ from the same laboratory has been extensively experimentally verified and a high success rate (67 %) was reported. We initially tested our approach on the design of the “language model design”, however, we observed that over-optimization of the designed proteins leads to low diversity *α*-helix bundles. The authors of this engine report a similar finding for a suboptimal setup of the annealing scheme.

We demonstrate here that parallel tempering can be applied in protein design. We must acknowledge that the current development of machine learning programs for protein design leads to very fast tools such as Protein Message Passing Neural Network ^7^ or ProteinGenerator ^15^. These tools can design stable proteins in very short time. In many applications, we can expect that these tools can outperform our approach, i.e. that a series of independent Protein Message Passing Neural Network or ProteinGenerator runs may be more efficient than our parallel tempering design.

However, we see great potential in the combination of parallel tempering protein design with some other physics-based approaches developed to improve sampling in molecular simulations or other applications ^16^. We will follow this path in the future.

## 4 METHODS

ESMfold was obtained from the GitHub repository (github.com/facebookresearch/esm). We used the code of “protein design programming language” ^3^. This code starts from a random amino acid sequence. The user can define key structure descriptors such as predicted Local Distance Difference Test (pLDDT), predicted Template Modeling (pTM), number of nonpolar surface residues, root mean square deviation from a target structure, symmetry descriptors and other variables. These variables are combined into a score (a lower score means higher agreement with desired properties). It uses a Monte Carlo simulated annealing to optimize the amino acid sequence.

The code was modified to provide start from a user-supplied amino acid sequence and to improve monitoring of the evolution of amino acid sequences. Parallel tempering was performed using a Python script.

The proteins were designed to maximize pLDDT, maximize pTM, minimize the number of surface hydrophobic residues, and maximize globularity (with weights 1, 1, 1 and 0.1, respectively). We performed design of proteins with 100 and 200 amino acid residues. The initial sequences were generated randomly. The temperatures were selected using a trial and error approach. We used 17 replicas (temperatures 0.00984, 0.0124, 0.0156, 0.0197, 0.0248, 0.0313, 0.0394, 0.0496, 0.0625, 0.0787, 0.099, 0.125, 0.157, 0.198, 0.25, 0.5 and 1.0, in *kT* units) for proteins with 100 residues. We used 24 replicas (temperatures 0.00164, 0.00195, 0.00232, 0.00276, 0.00328, 0.00391, 0.00465, 0.00552, 0.00657, 0.00781, 0.00929, 0.0110, 0.0131, 0.0156, 0.0221, 0.0313, 0.0442, 0.0625, 0.0884, 0.125, 0.177, 0.25, 0.5 and 1.0, in *kT* units) for proteins with 200 residues. Replica exchange attempts were made every 100 Monte Carlo steps, unless otherwise stated. There were exchange attempts between replicas 0 and 1, 2 and 3 etc. in odd exchange attempts and between replicas 1 and 2, 3 and 4 etc. in even exchange attempts. Due to restriction in available hardware, we performed design campaigns as jobs consisting of 25 or 10 exchange attempts, for proteins of 100 and 200 residues, respectively. There were no replica exchanges between the end of one job and the beginning of the subsequent job.

A standard Monte Carlo simulated annealing was performed on selected structures (Figure 2E and Figure 3E) to improve their stability. This consisted of 1,000 steps. The initial temperature was set to the temperature at which the design was sampled. The temperature was reduced by a factor of 0.97 after each step.

For visualizations, we used ESMfold ^11^, Python, VMD ^17^, and R. All material (codes, raw data, visualization codes, etc.) can be obtained at Zenodo (DOI: 10.5281/zenodo.15005147).

## 5 CONCLUSIONS

We extend the parallel tempering algorithm to protein design. It can be applied as an extension of methods commonly used in protein design, namely Monte Carlo sampling and simulation annealing. Parallel tempering shows the potential to optimize protein sequences more efficiently than conventional methods. Our approach was not experimentally validated by protein expression and characterization. However, we believe that the concept can be used in different protein design engines (the way energy of a protein sequence is calculated) and that the performance in experimental validation is most likely determined by the performance of the protein design engine. The advantage of our approach is the fact that it can provide a continuous flow of designed sequences.

## Abbreviations

ESM: evolutionary scale modeling
pLDDT: predicted local distance difference test
pTM: predicted template modeling
VMD: visual molecular dynamics.

## AUTHOR CONTRIBUTIONS

**Preet Kalani**: Methodology; software; investigation; writing

- review and editing; visualization. **Vojtěch Spiwok**: Conceptualization; methodology; software; investigation; writing
- original draft; writing – review and editing; visualization; data curation; resources; supervision; project administration; funding acquisition.

## ACKNOWLEDGMENTS

This work was supported by COST (project ML4NGP, CA21160). Participation in the COST project was supported by the Ministry of Education, Youth and Sports of the Czech Republic (LUC 24136). Long-term availability of the resulting tools and data is supported by ELIXIR CZ (LM 2023055).

## CONFLICT OF INTEREST

The authors declare no potential conflict of interests.

## SUPPORTING INFORMATION

Additional supporting information can be found online in the Supporting Information section at the end of this article.

## Notes

This research was supported by the COST and the Ministry of Education, Youth and Sports of Czech Republic (ML4NGP, CA21160, LUC 24136, LM 2023055)

https://doi.org/10.5281/zenodo.15005147

### Competing Interest Statement

The authors have declared no competing interest.

