## Supporting Information for "Design of proteins by parallel tempering in the sequence space"

**Preet Kalani** | **Vojtěch Spiwok**

[1]Department of Biochemistry and Microbiology, University of Chemistry and Technology, Prague, Technická 3, Prague 6, 166 28, Czech Republic

**Correspondence**
Corresponding author Vojtěch Spiwok.


### Table of Contents

**Figure S1-S17**: Demultiplexed replicas of design of 100-residue proteins by parallel tempering.

**Figure S18**: Design of 100-residue proteins by parallel tempering with a higher replica exchange frequency.

**Figure S19-S42**: Demultiplexed replicas of design of 200-residue proteins by parallel tempering.

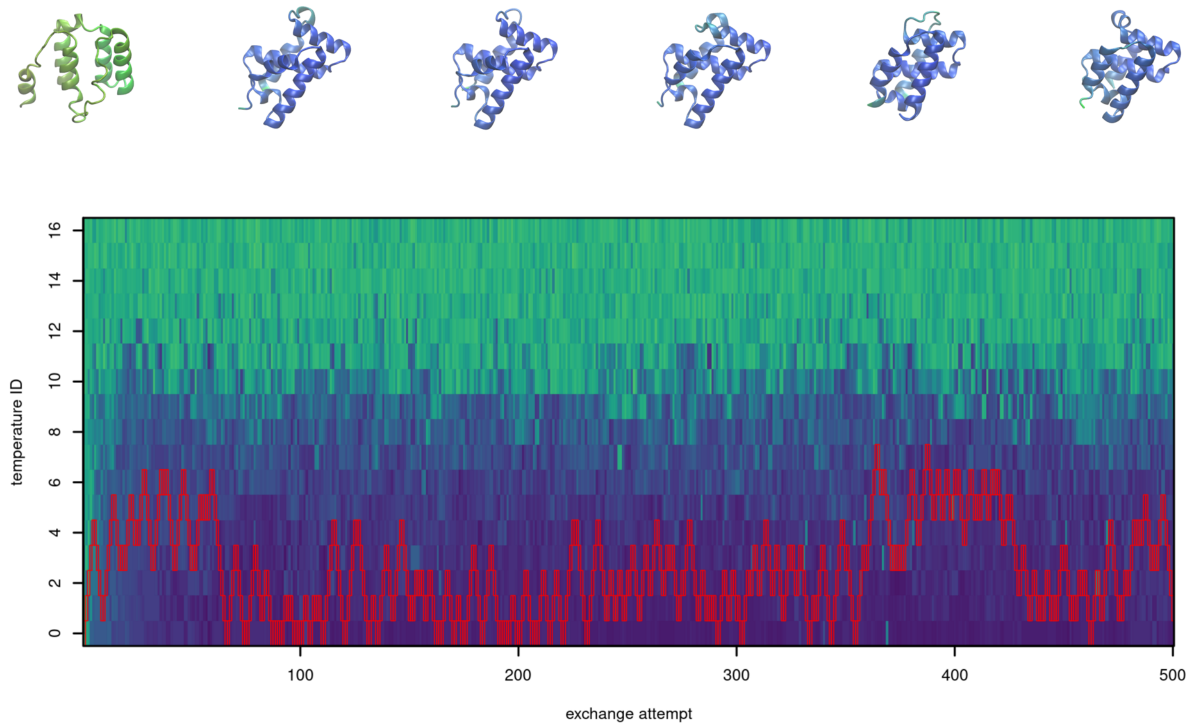


**FIGURE S1** Demultiplexed replica 0 of design of 100-residue proteins by parallel tempering.

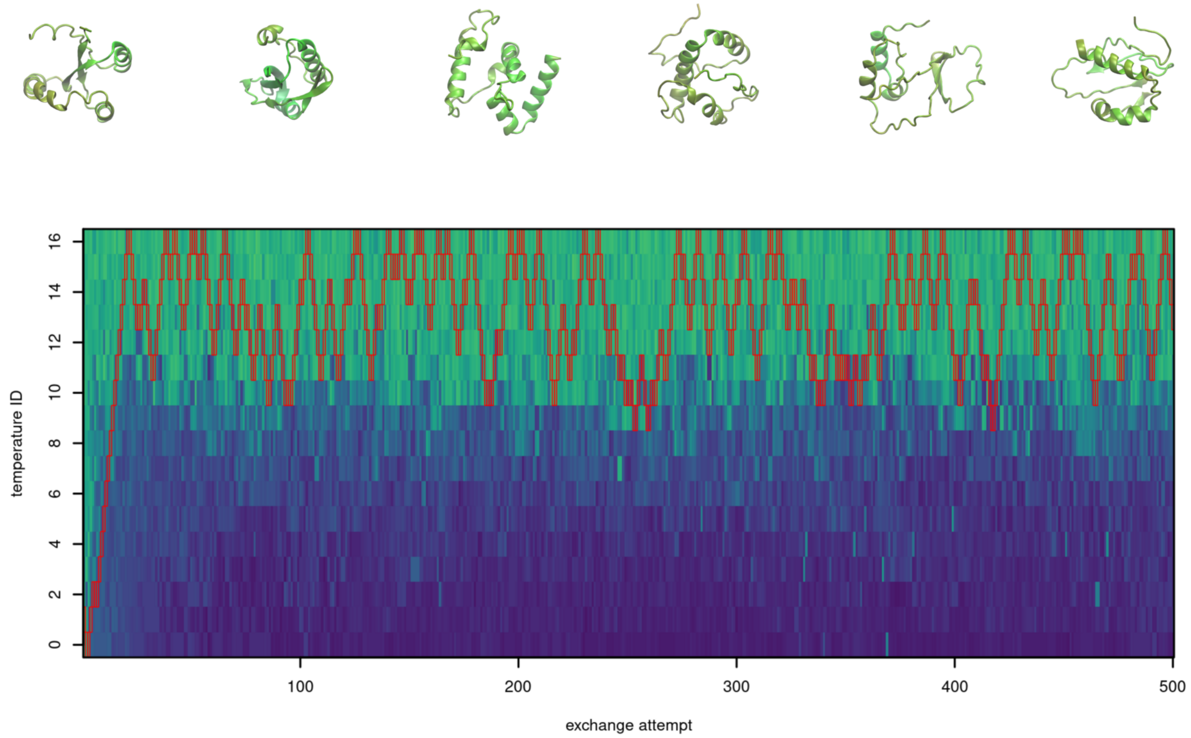


**FIGURE S2** Demultiplexed replica 1 of design of 100-residue proteins by parallel tempering.

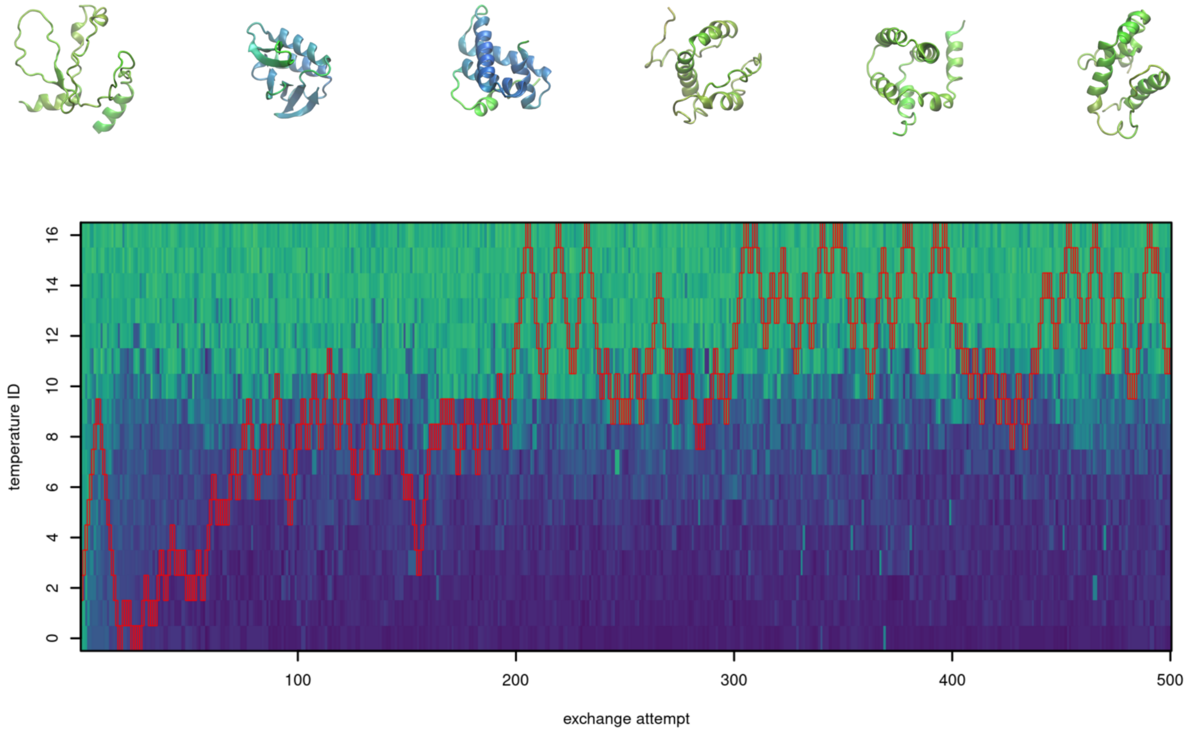


**FIGURE S3** Demultiplexed replica 2 of design of 100-residue proteins by parallel tempering.

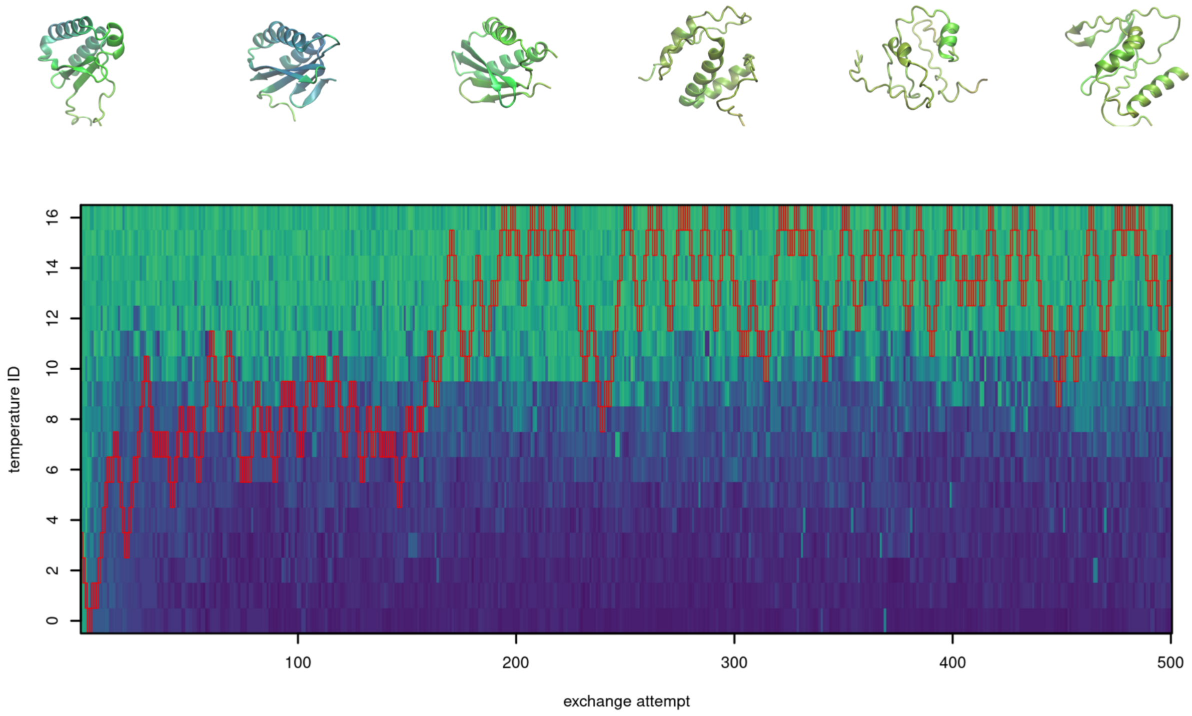


**FIGURE S4** Demultiplexed replica 3 of design of 100-residue proteins by parallel tempering.

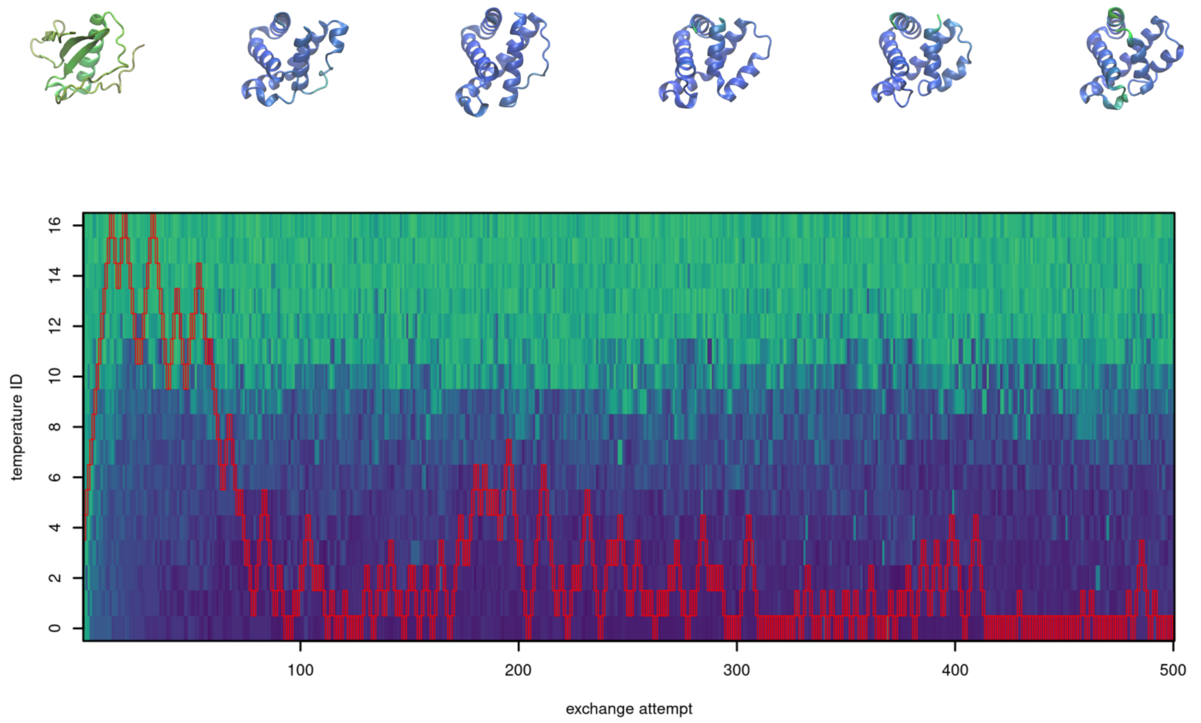


**FIGURE S5** Demultiplexed replica 4 of design of 100-residue proteins by parallel tempering.

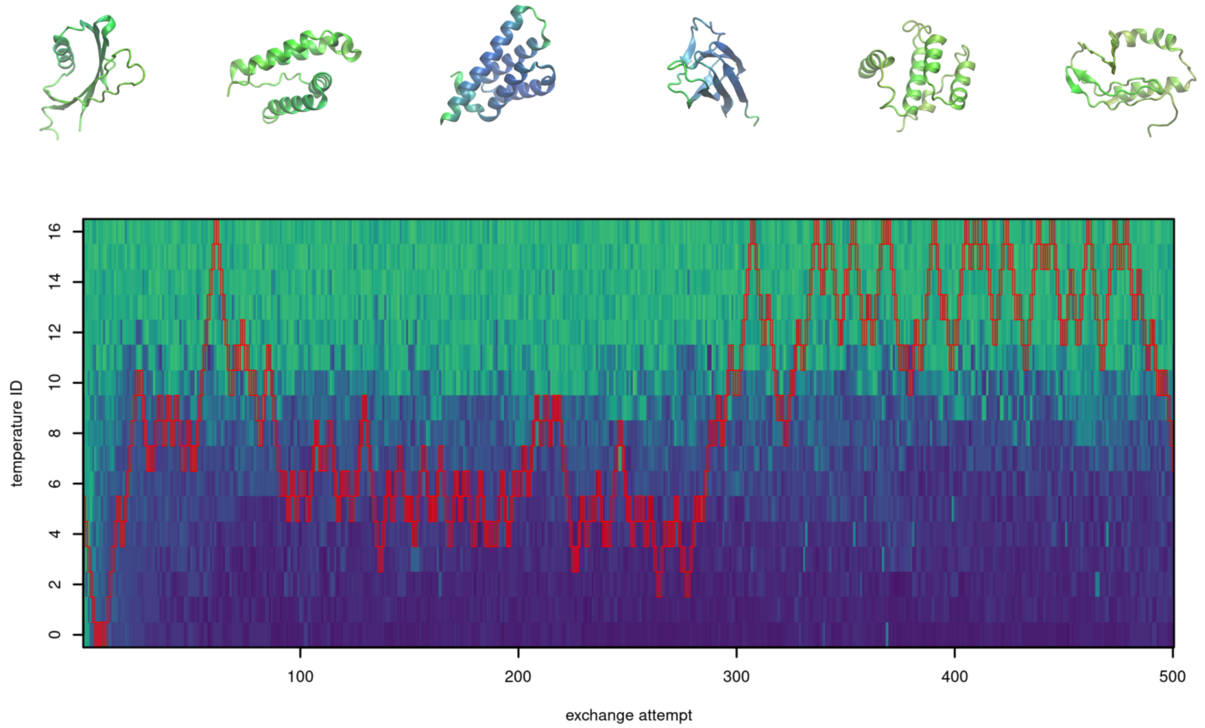


**FIGURE S6** Demultiplexed replica 5 of design of 100-residue proteins by parallel tempering.

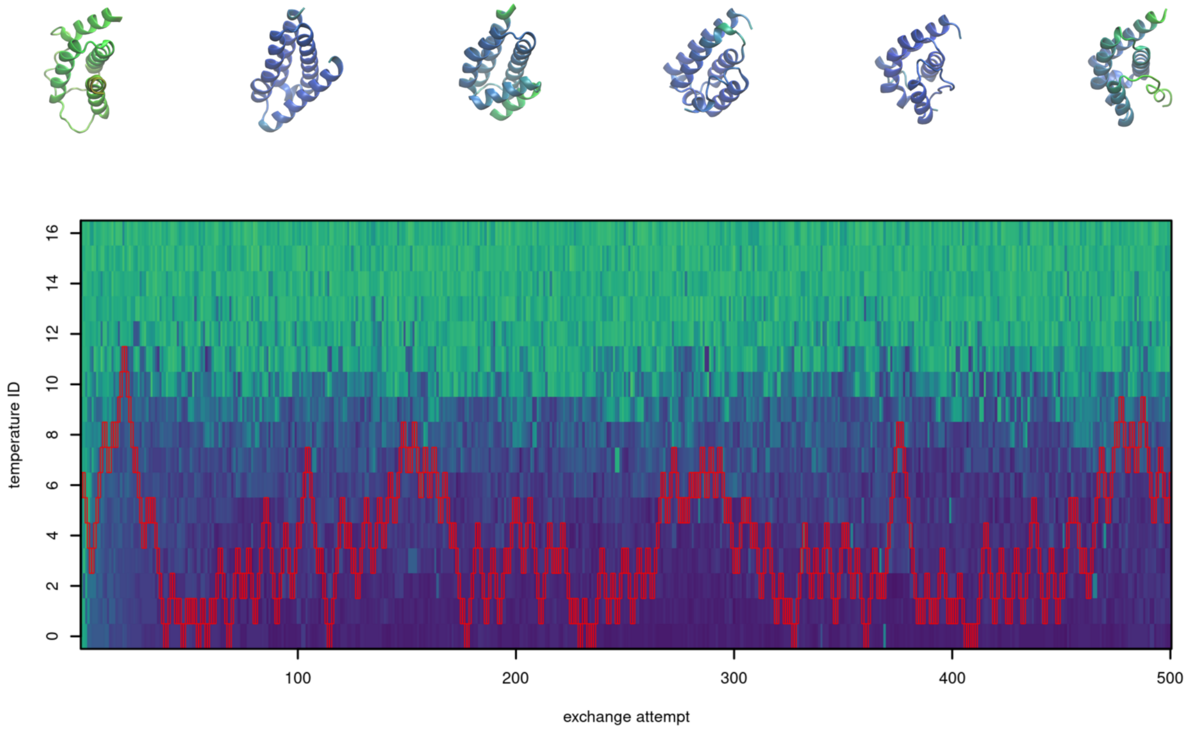


**FIGURE S7** Demultiplexed replica 6 of design of 100-residue proteins by parallel tempering.

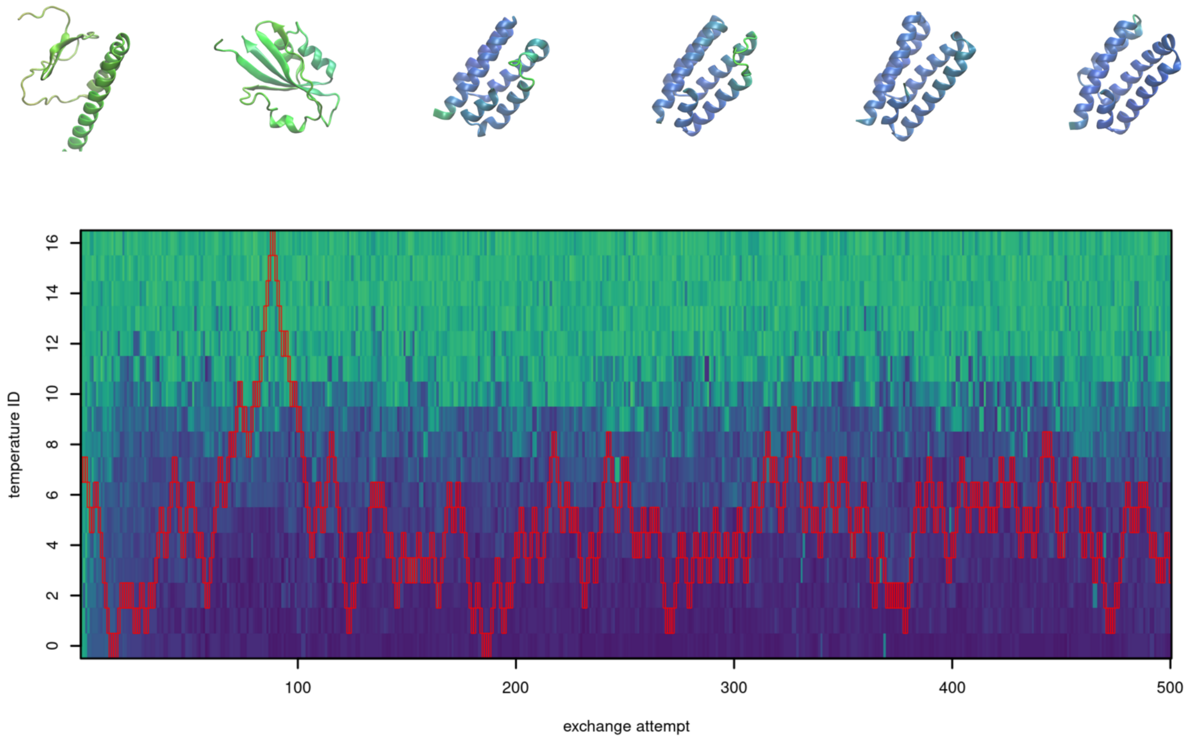


**FIGURE S8** Demultiplexed replica 7 of design of 100-residue proteins by parallel tempering.

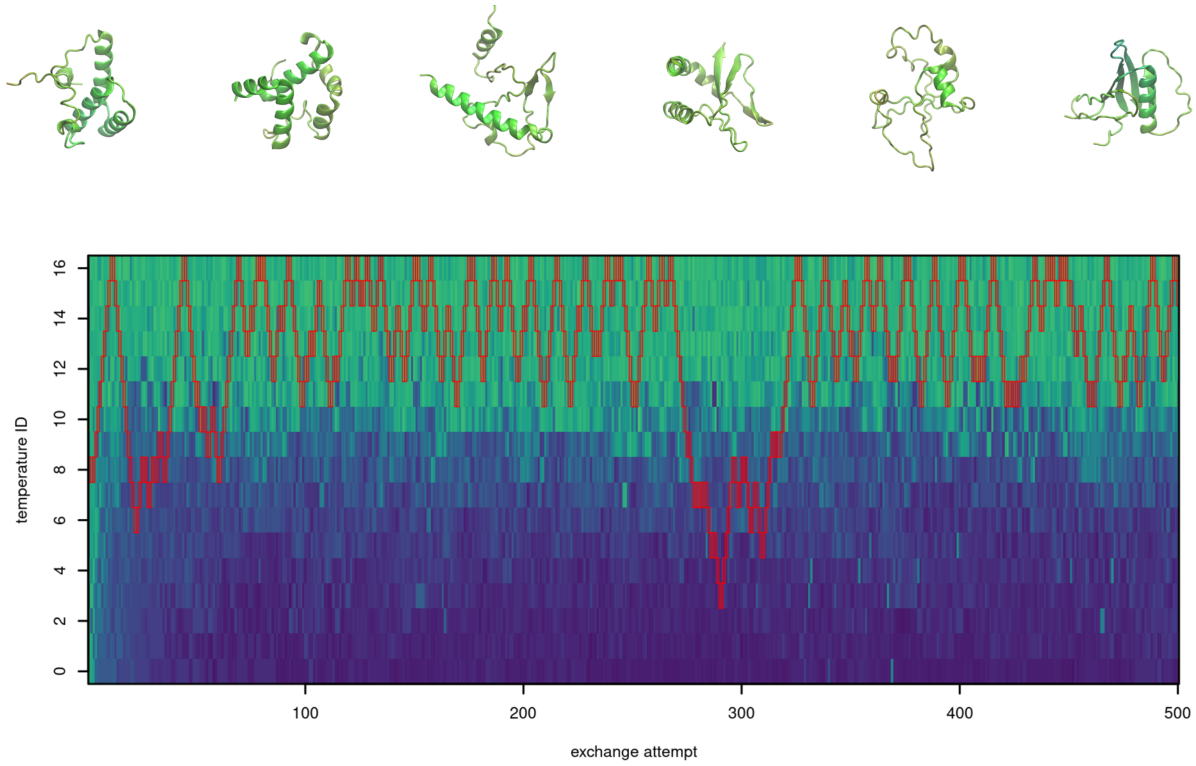


**FIGURE S9** Demultiplexed replica 8 of design of 100-residue proteins by parallel tempering.

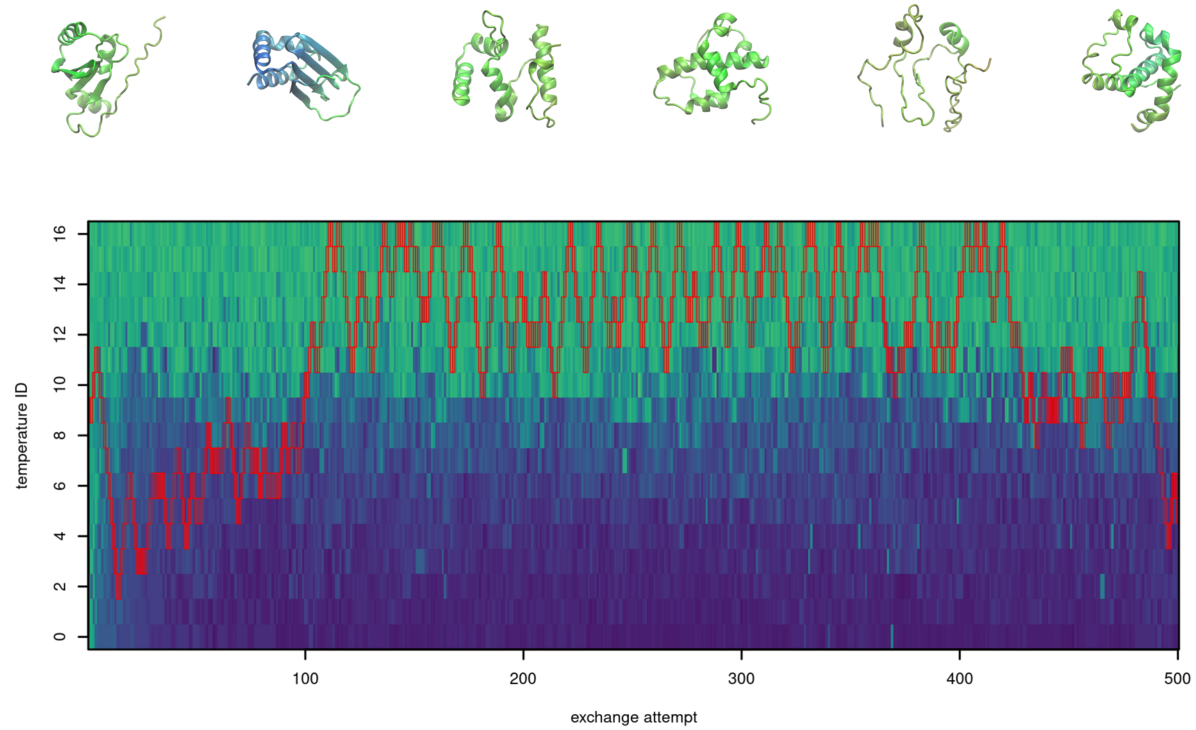


**FIGURE S10** Demultiplexed replica 9 of design of 100-residue proteins by parallel tempering.

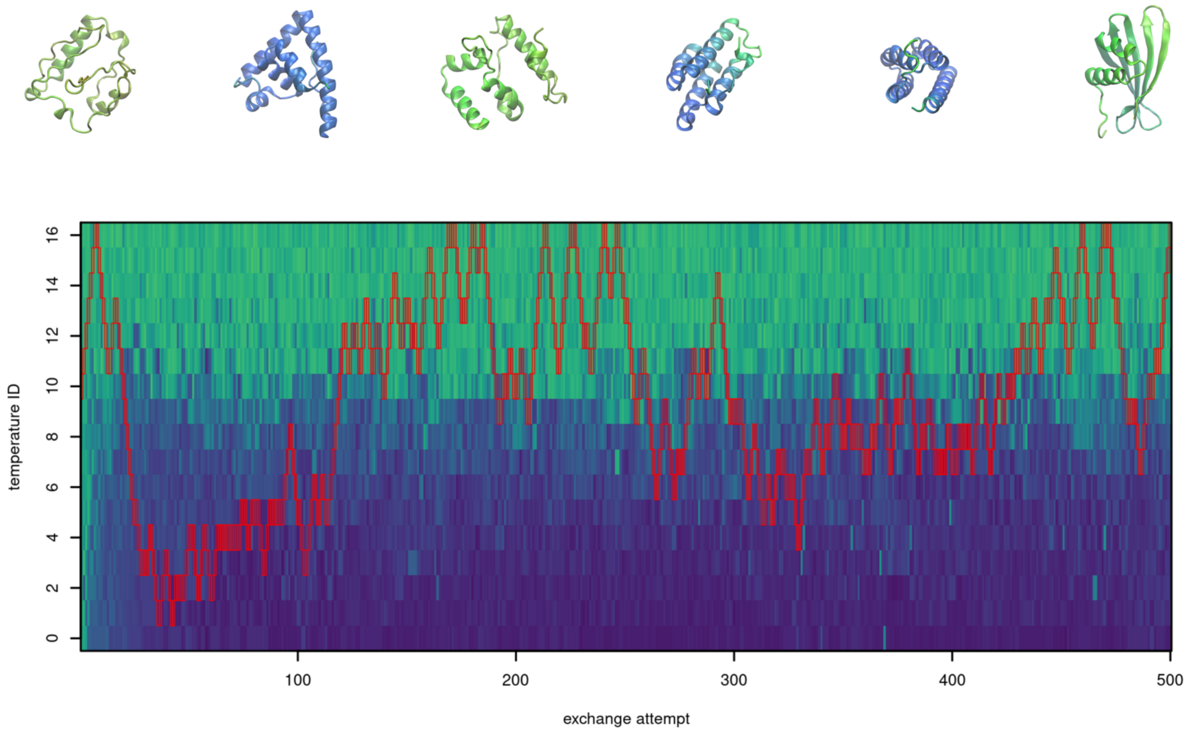


**FIGURE S11** Demultiplexed replica 10 of design of 100-residue proteins by parallel tempering.

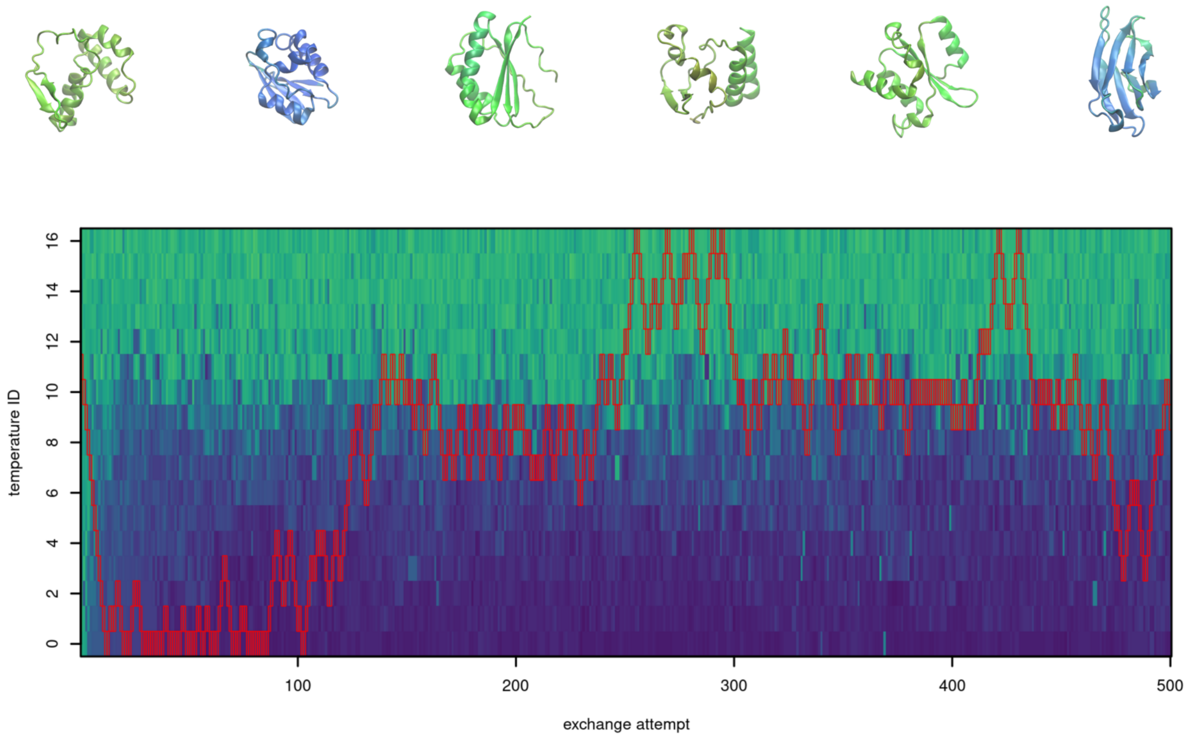


**FIGURE S12** Demultiplexed replica 11 of design of 100-residue proteins by parallel tempering.

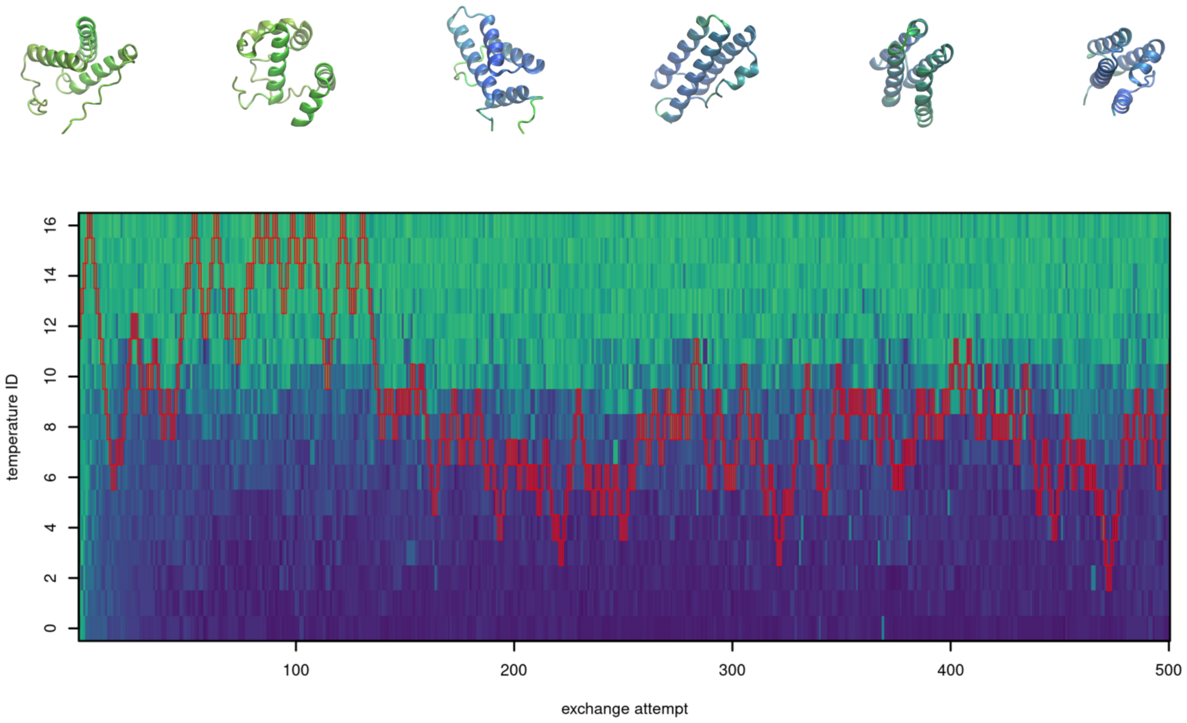


**FIGURE S13** Demultiplexed replica 12 of design of 100-residue proteins by parallel tempering.

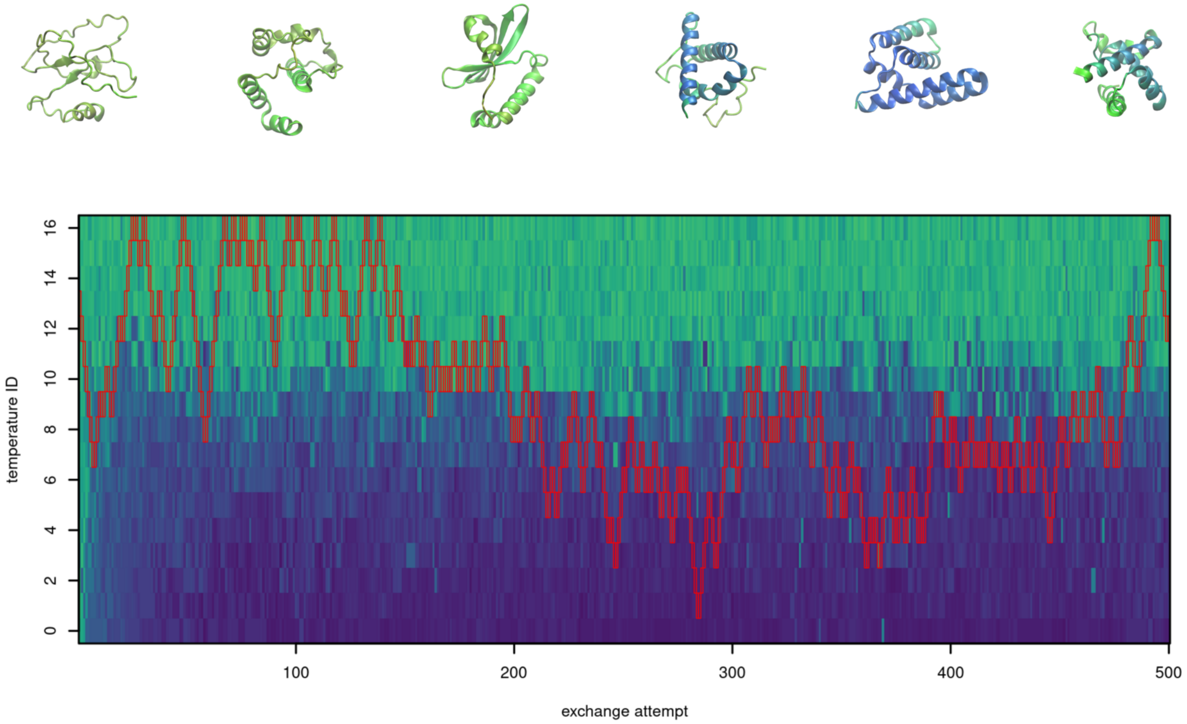


**FIGURE S14** Demultiplexed replica 13 of design of 100-residue proteins by parallel tempering.

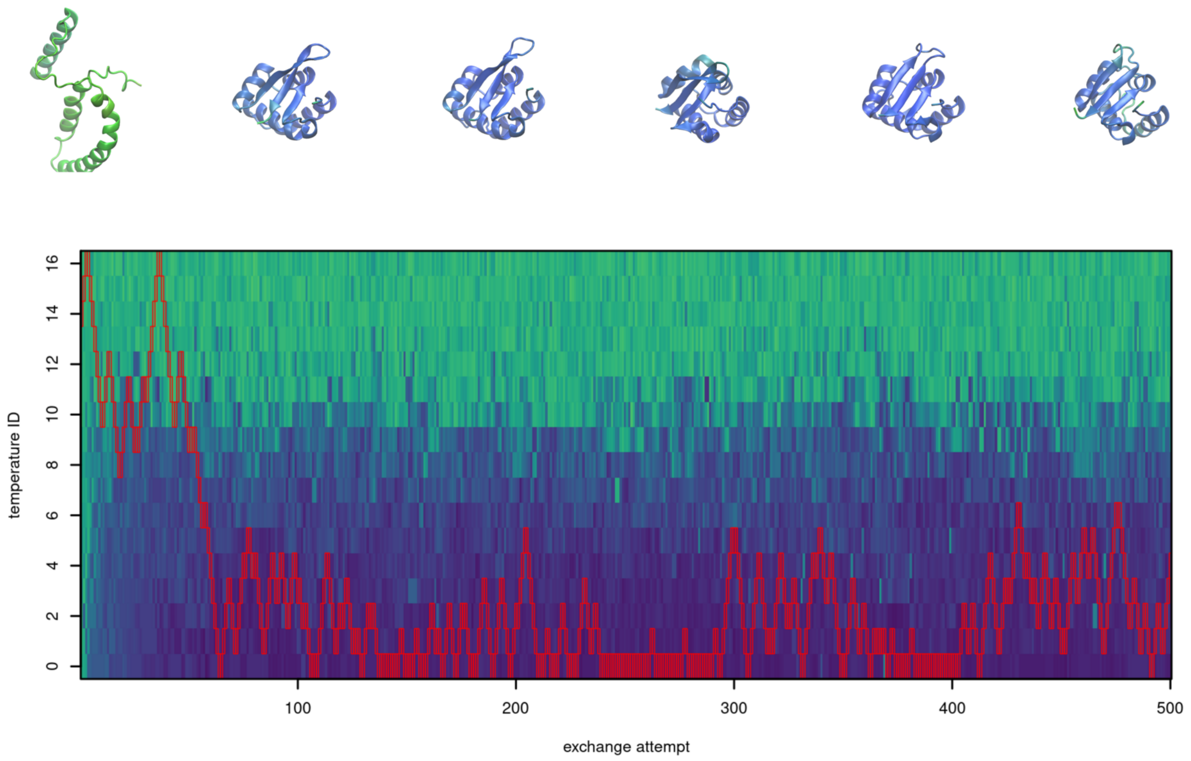


**FIGURE S15** Demultiplexed replica 14 of design of 100-residue proteins by parallel tempering.

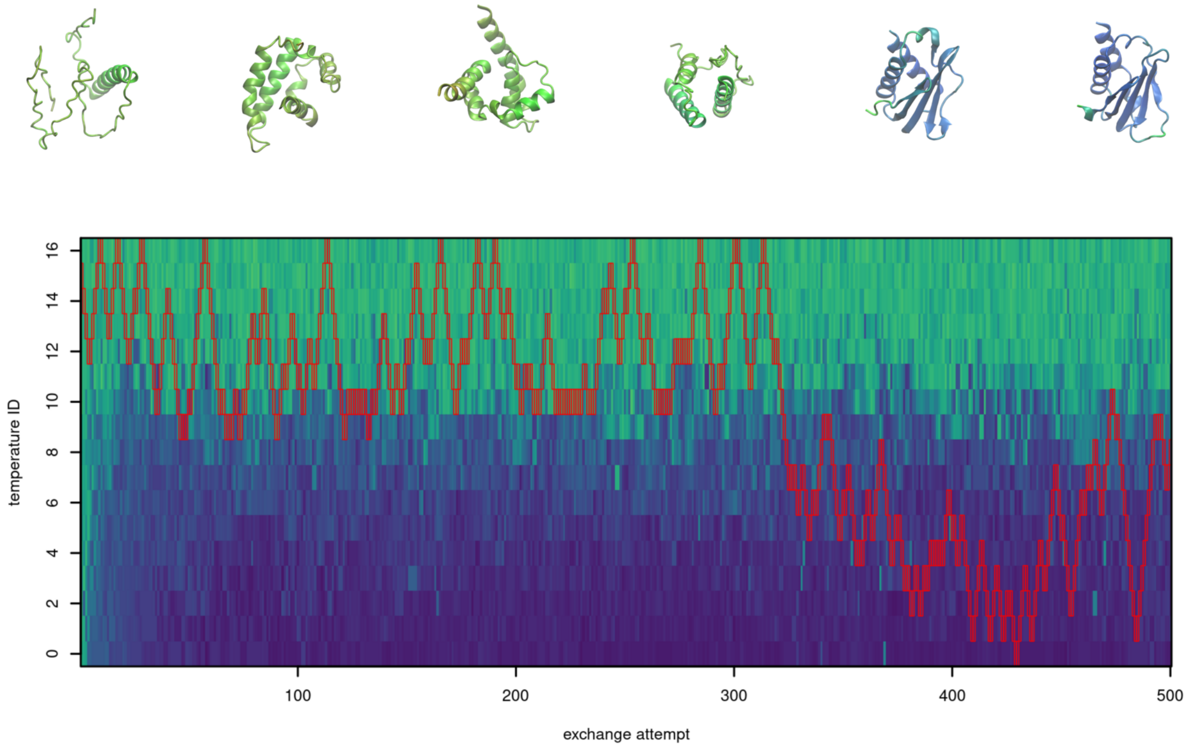


**FIGURE S16** Demultiplexed replica 15 of design of 100-residue proteins by parallel tempering.

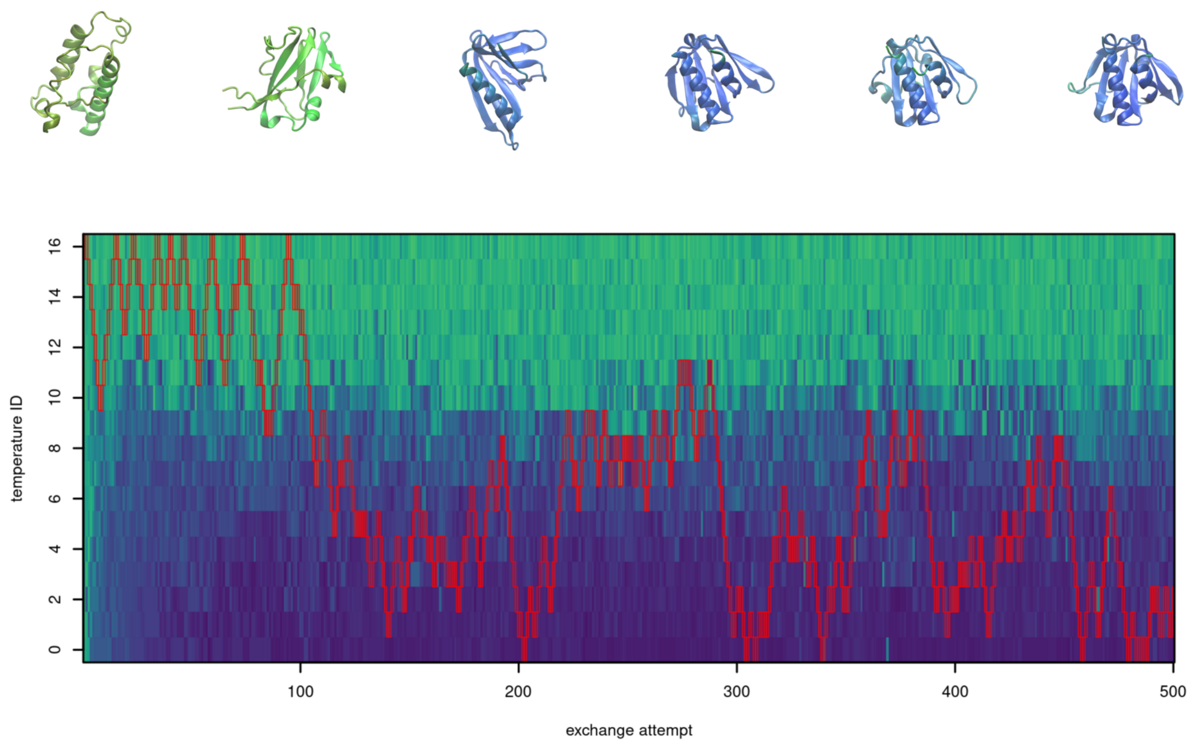


**FIGURE S17** Demultiplexed replica 16 of design of 100-residue proteins by parallel tempering.

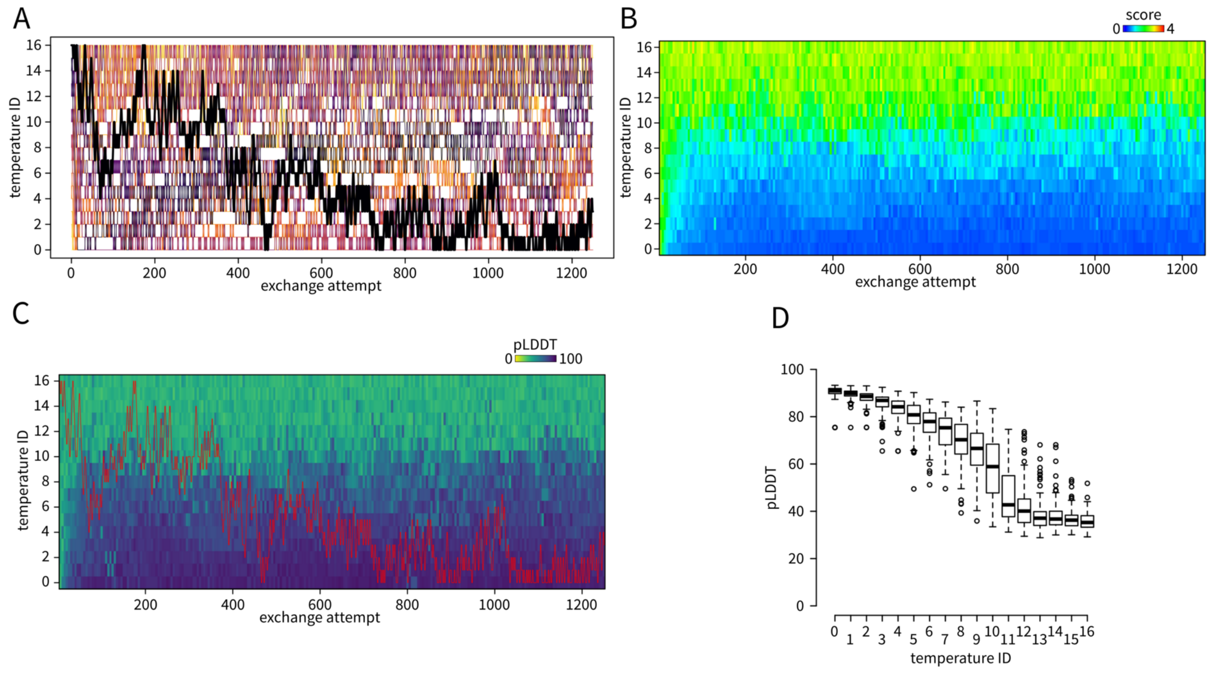


**FIGURE S18** Design of 100-residue proteins by parallel tempering with a higher replica exchange frequency. A – replica exchanges. Selected replica starting from temperature ID 16 is highlighted in tick. B – evolution of the score. C – evolution of mean C$\alpha$-pLDDT. The replica starting from temperature ID 16 is highlighted by red boxes. D – boxplot of the mean C$\alpha$-pLDDT at different temperatures (first 100 replica exchange attempts skipped, sampled at replica exchange points).

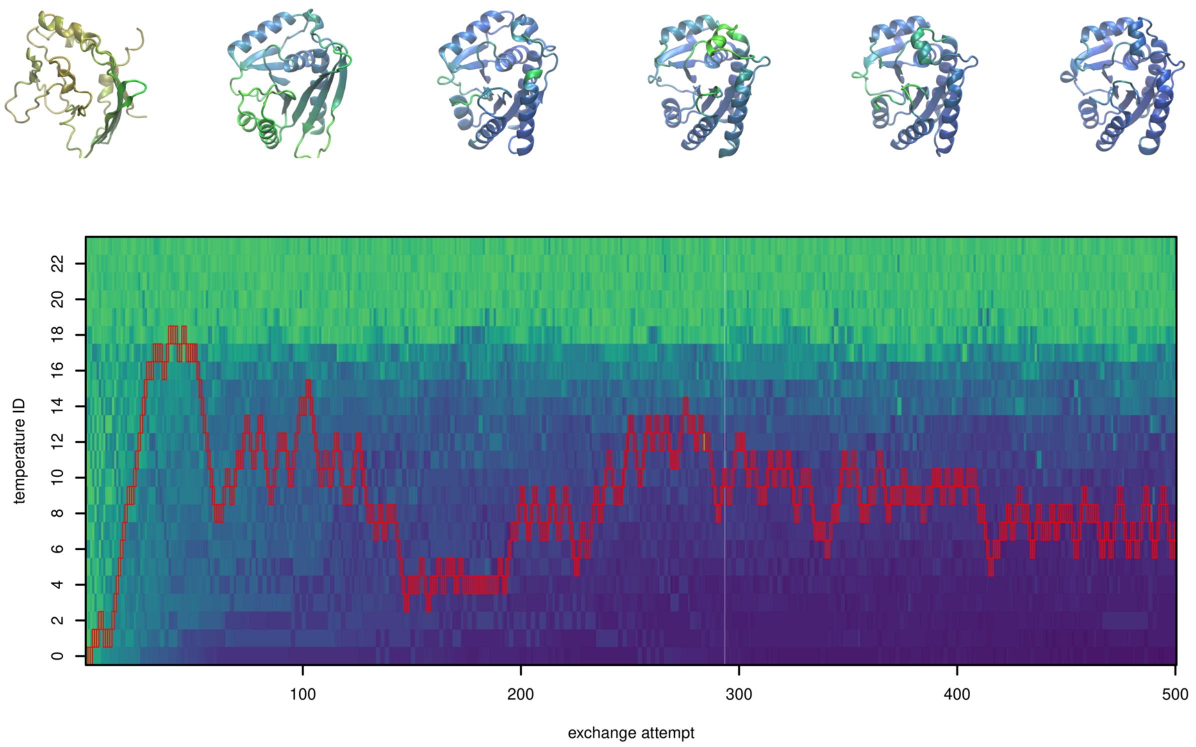


**FIGURE S19** Demultiplexed replica 0 of design of 200-residue proteins by parallel tempering.

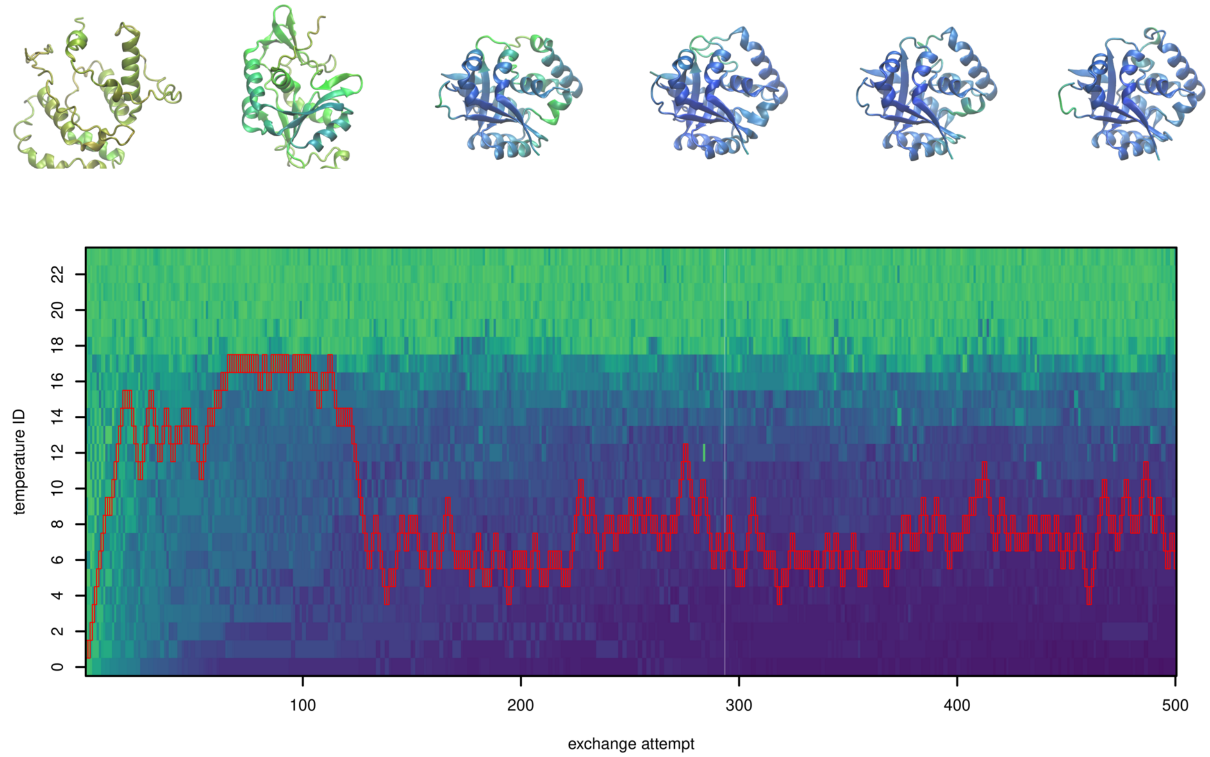


**FIGURE S20** Demultiplexed replica 1 of design of 200-residue proteins by parallel tempering.

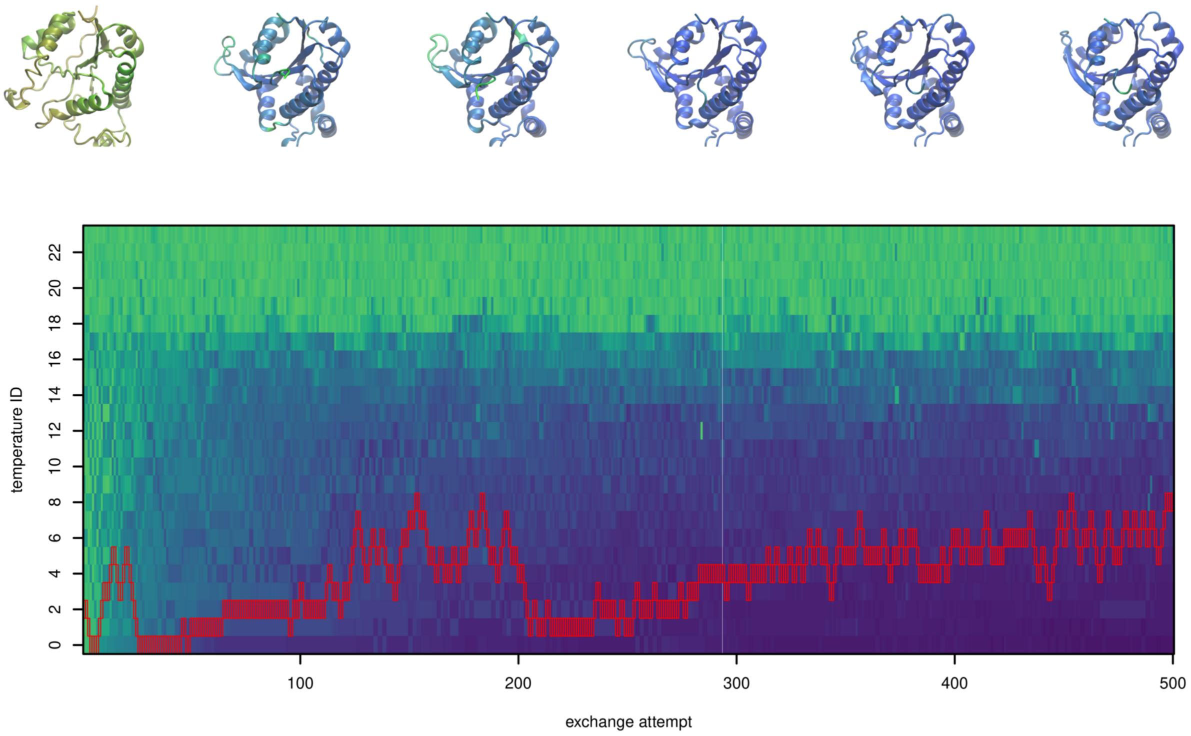


**FIGURE S21** Demultiplexed replica 2 of design of 200-residue proteins by parallel tempering.

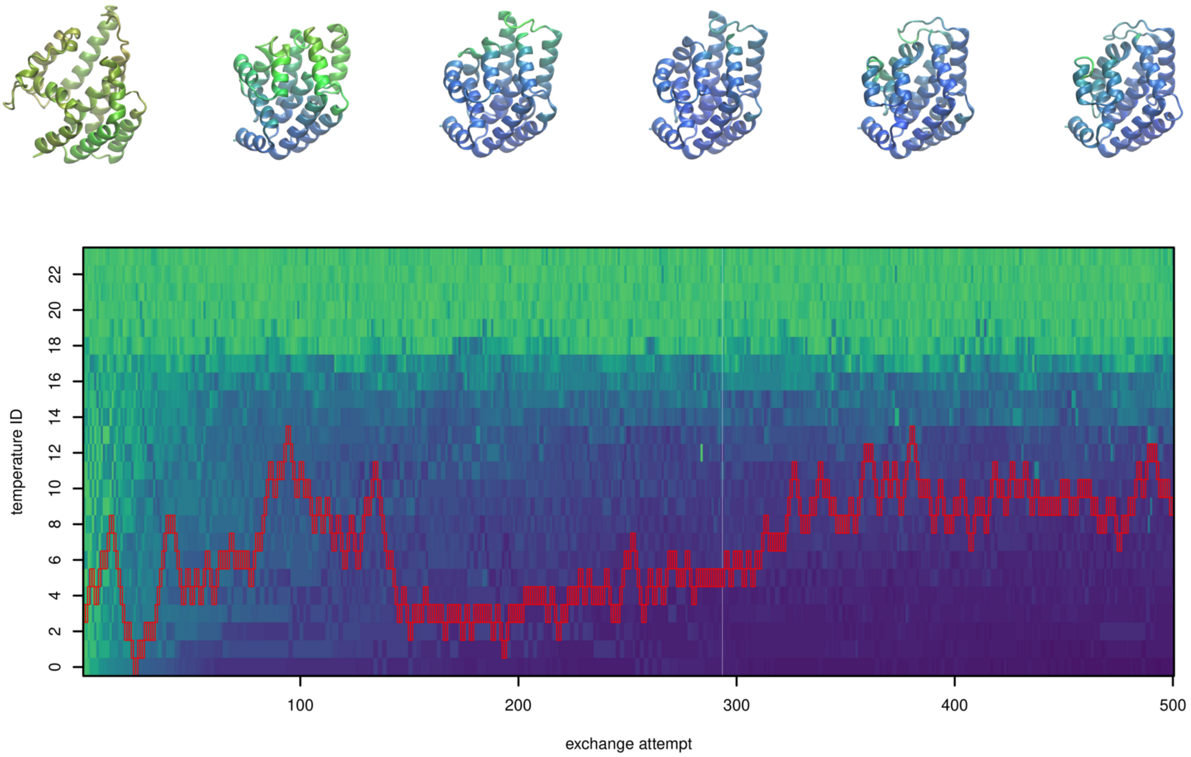


**FIGURE S22** Demultiplexed replica 3 of design of 200-residue proteins by parallel tempering.

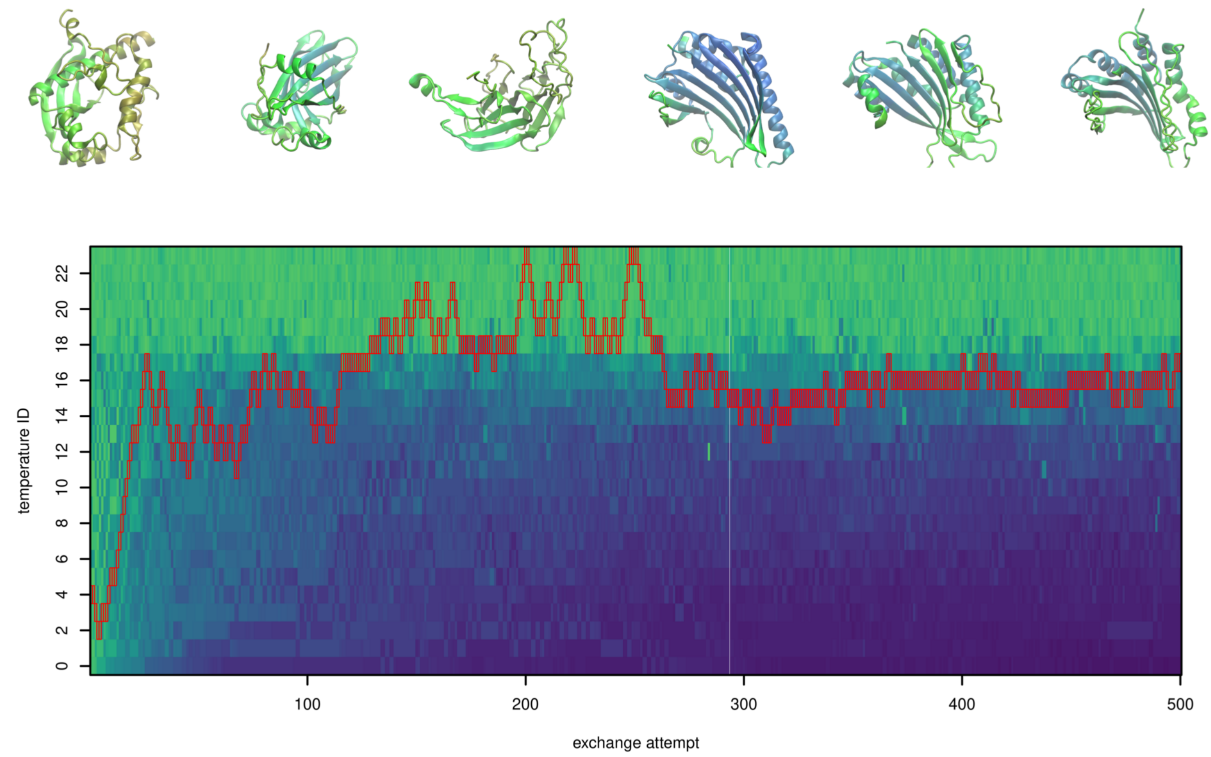


**FIGURE S23** Demultiplexed replica 4 of design of 200-residue proteins by parallel tempering.

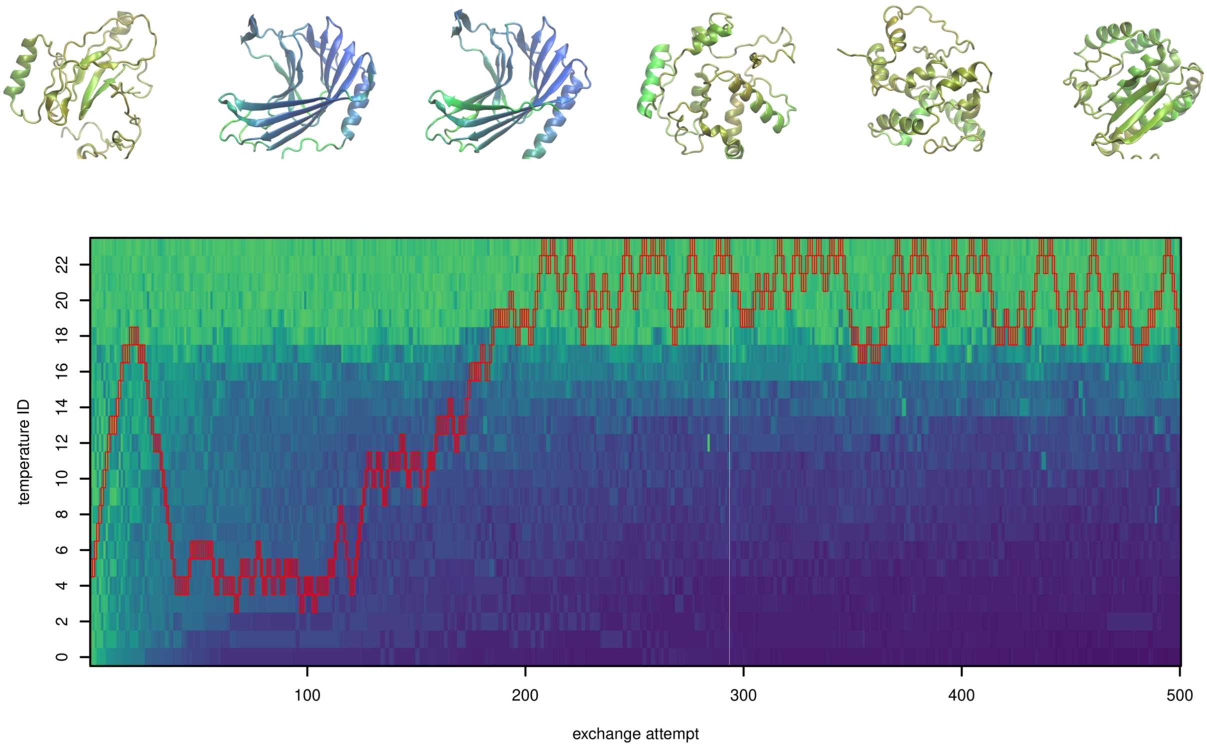


**FIGURE S24** Demultiplexed replica 5 of design of 200-residue proteins by parallel tempering.

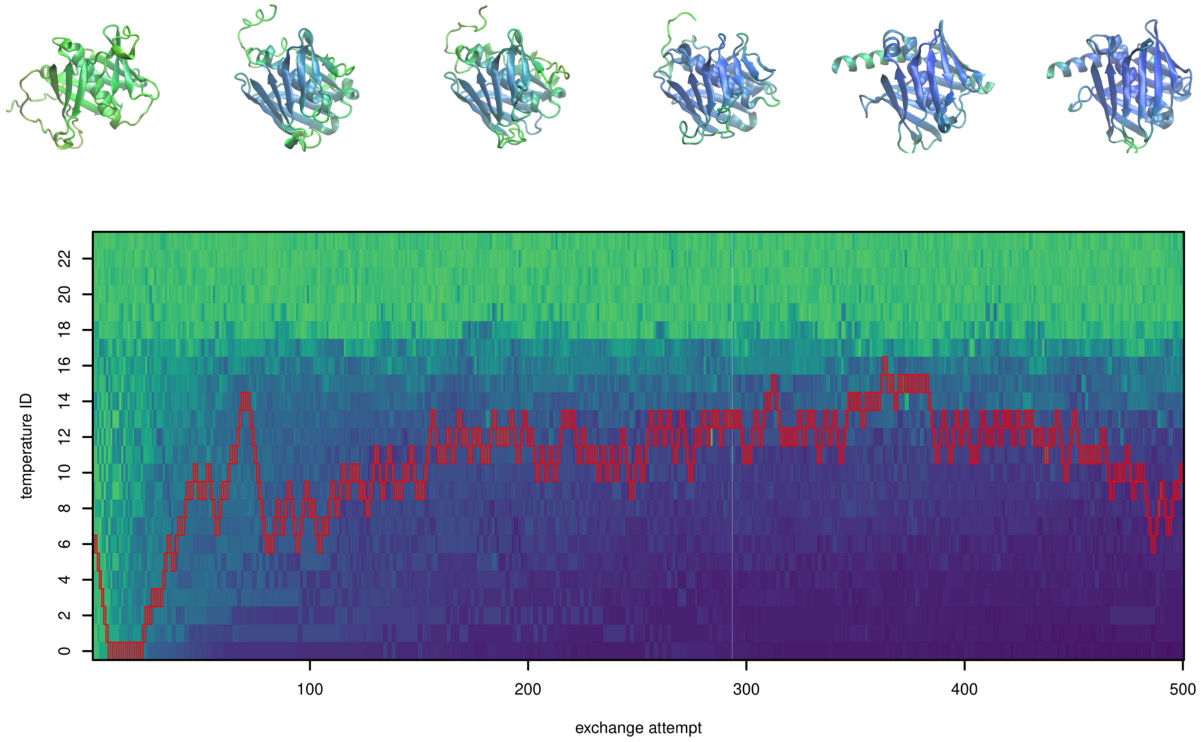


**FIGURE S25** Demultiplexed replica 6 of design of 200-residue proteins by parallel tempering.

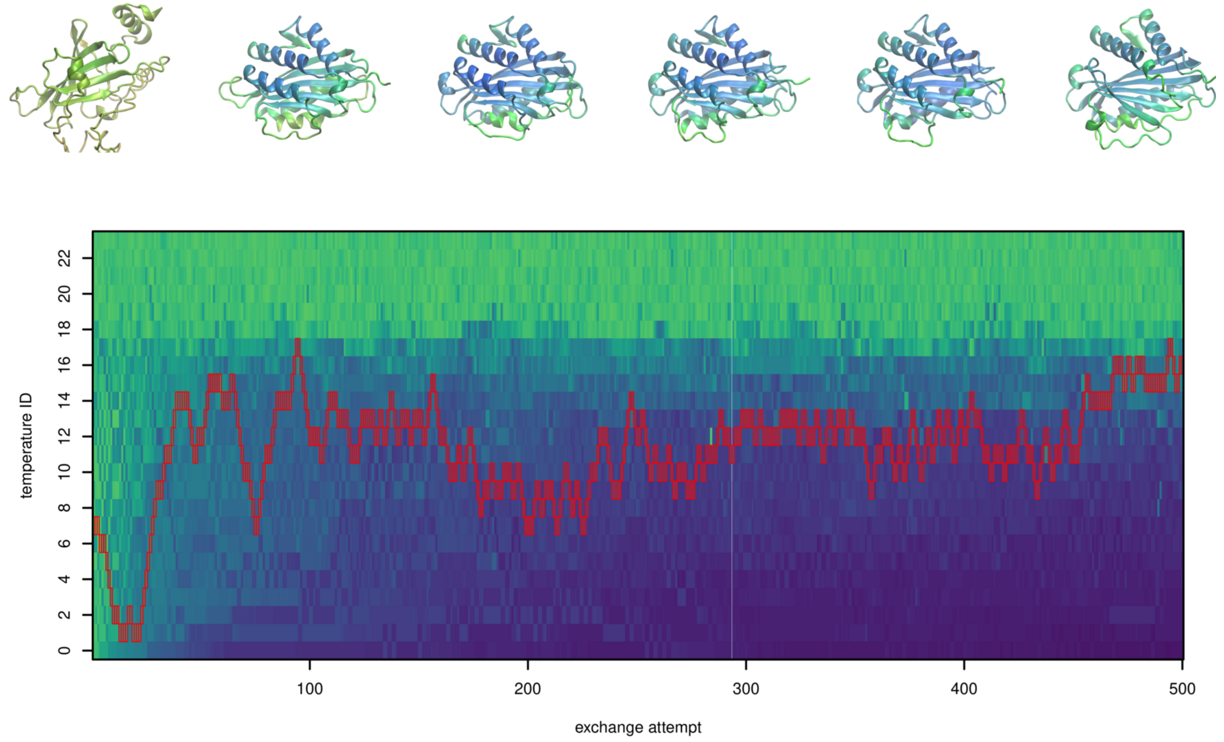


**FIGURE S26** Demultiplexed replica 7 of design of 200-residue proteins by parallel tempering.

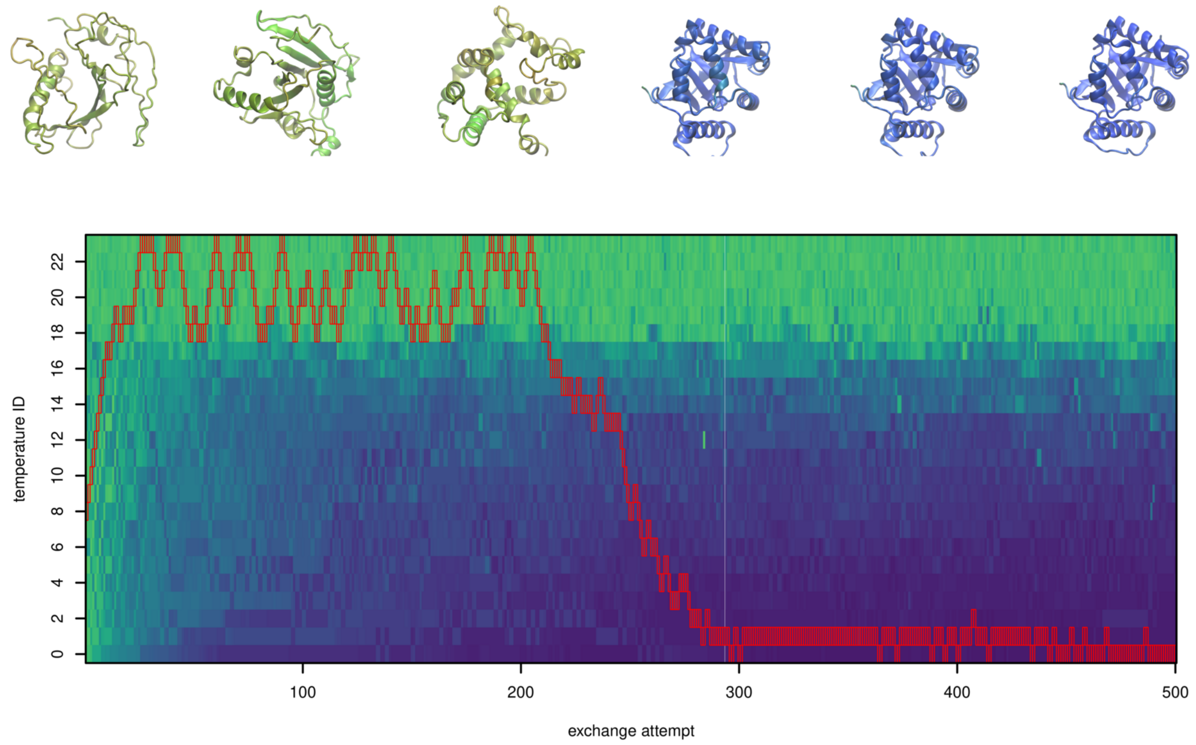


**FIGURE S27** Demultiplexed replica 8 of design of 200-residue proteins by parallel tempering.

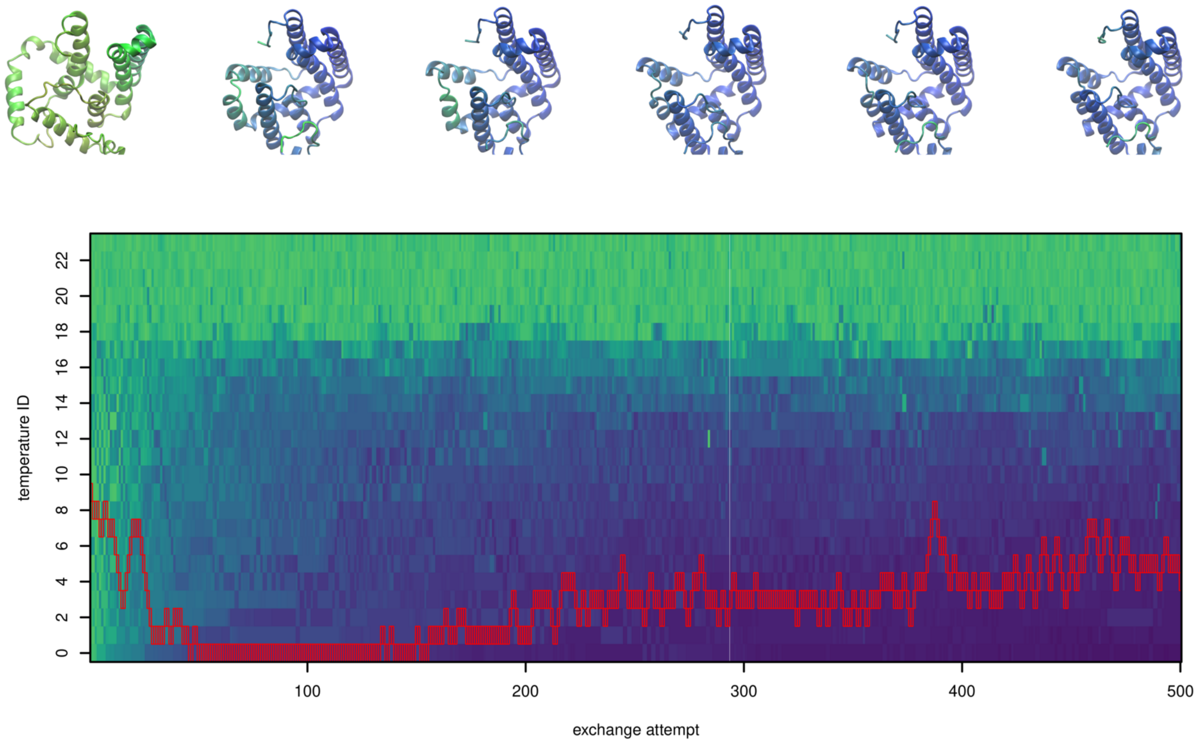


**FIGURE S28** Demultiplexed replica 9 of design of 200-residue proteins by parallel tempering.

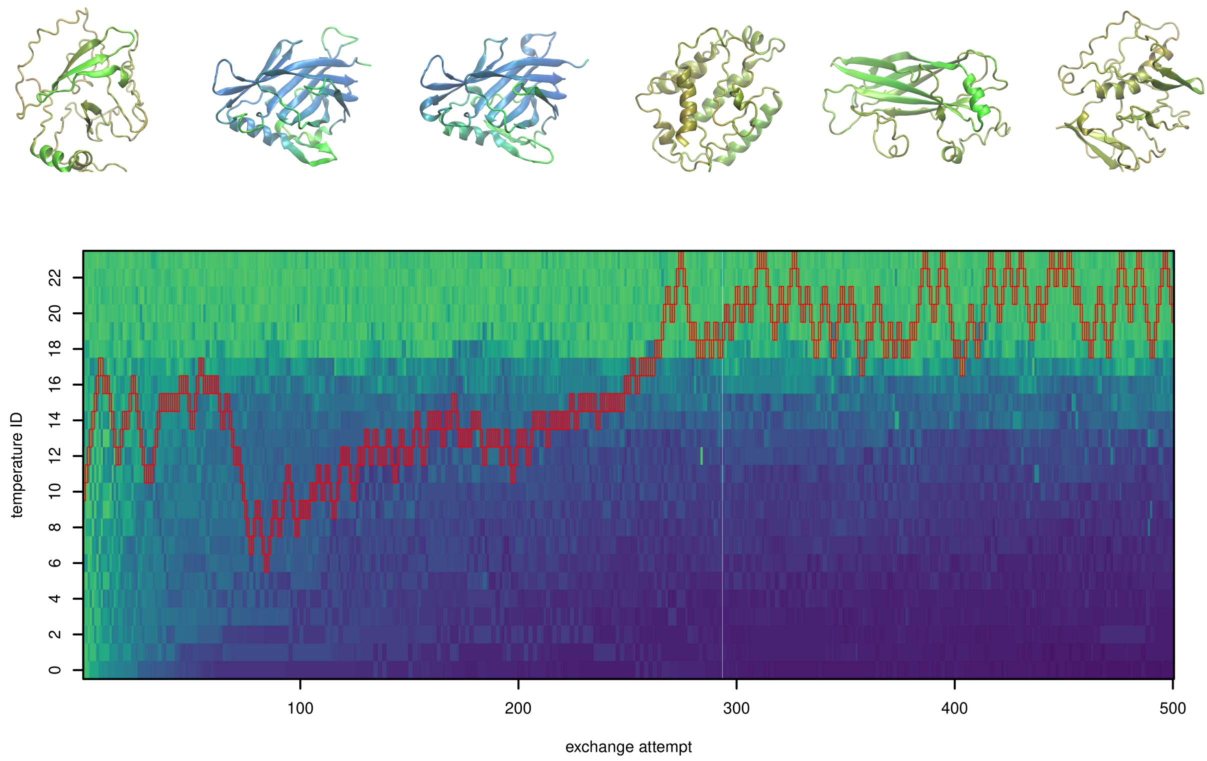


**FIGURE S29** Demultiplexed replica 10 of design of 200-residue proteins by parallel tempering.

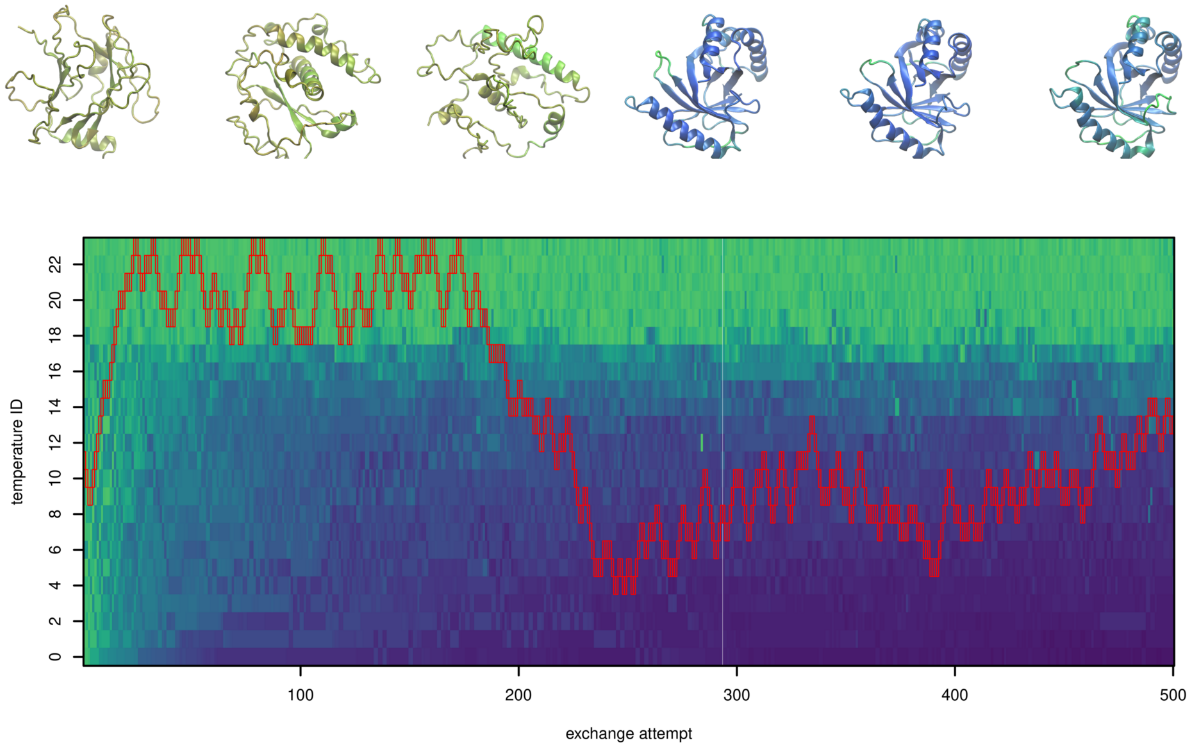


**FIGURE S30** Demultiplexed replica 11 of design of 200-residue proteins by parallel tempering.

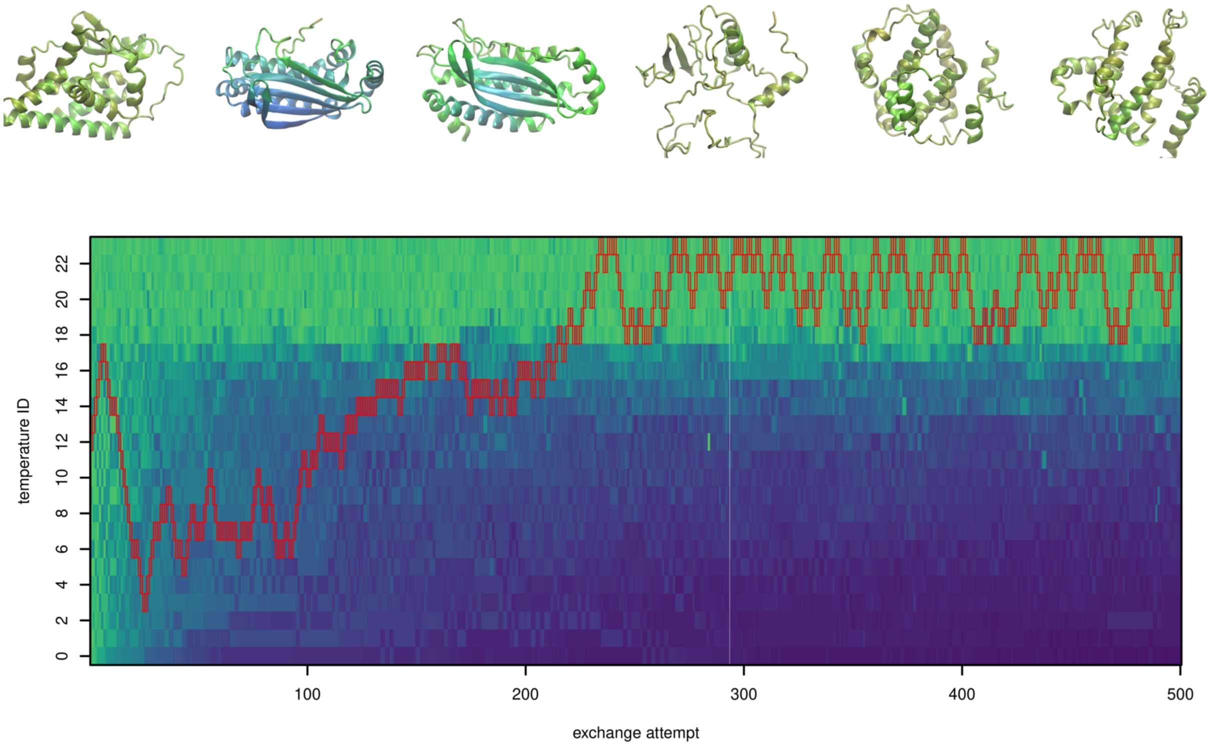


**FIGURE S31** Demultiplexed replica 12 of design of 200-residue proteins by parallel tempering.

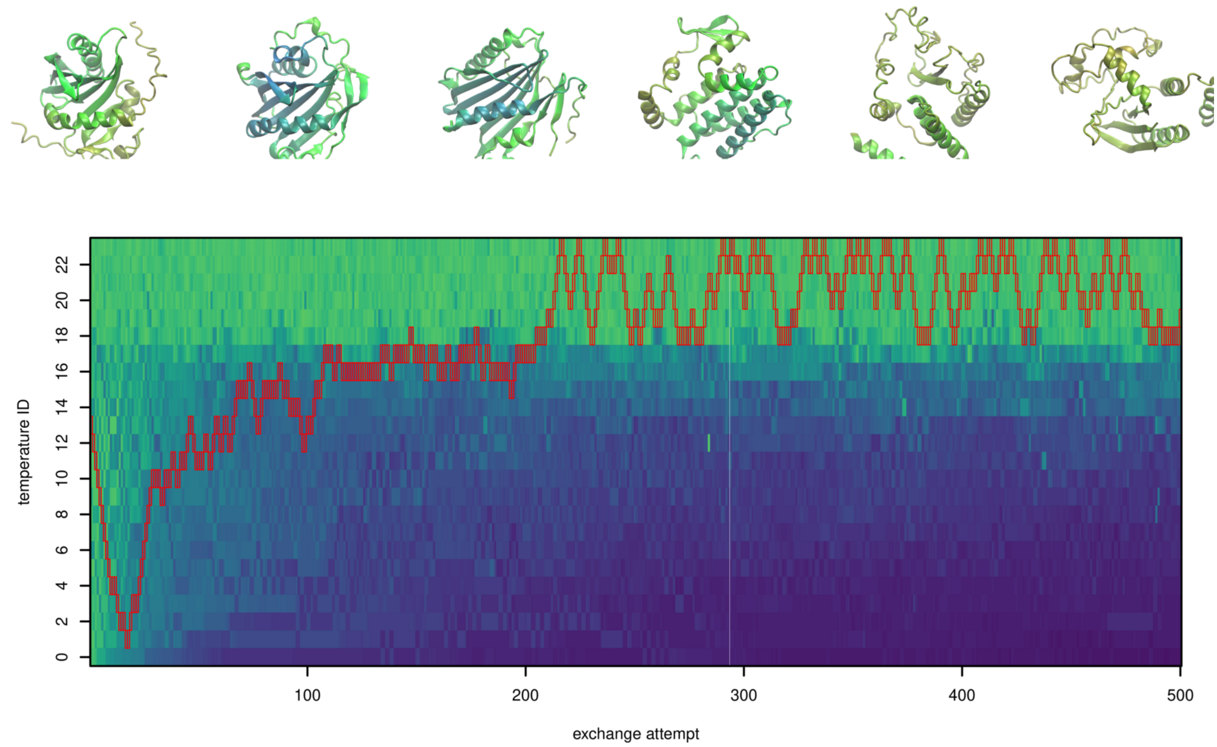


**FIGURE S32** Demultiplexed replica 13 of design of 200-residue proteins by parallel tempering.

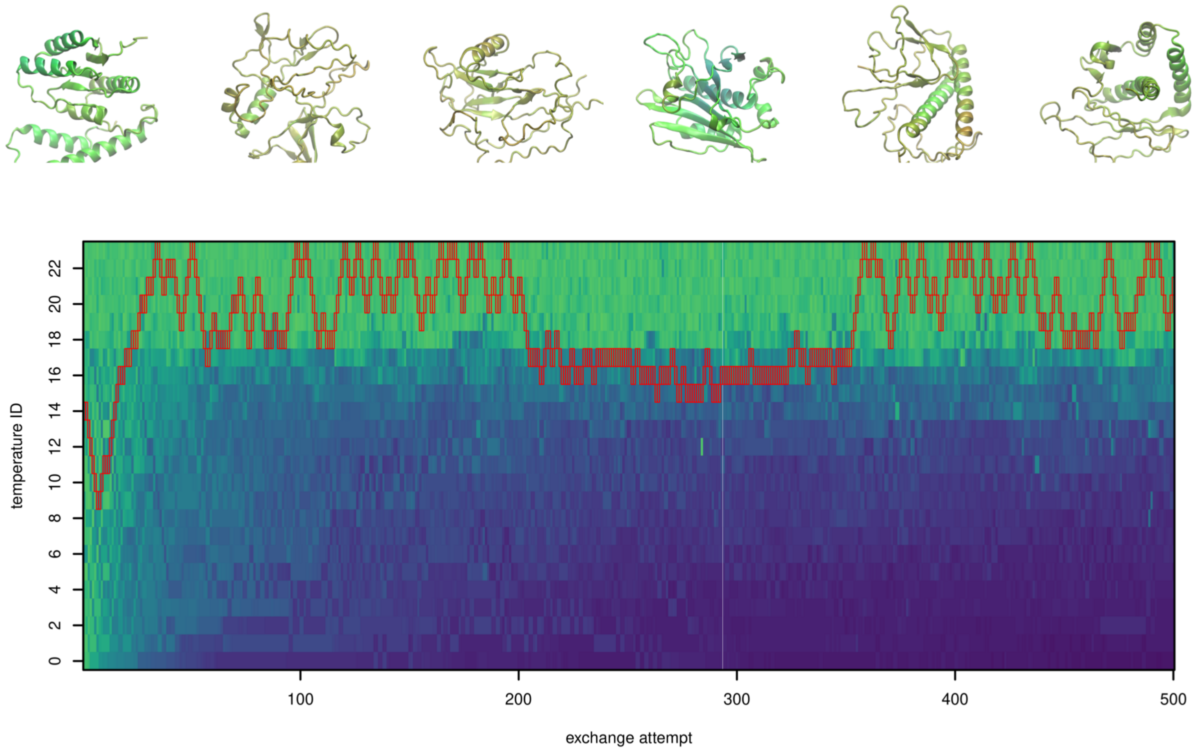


**FIGURE S33** Demultiplexed replica 14 of design of 200-residue proteins by parallel tempering.

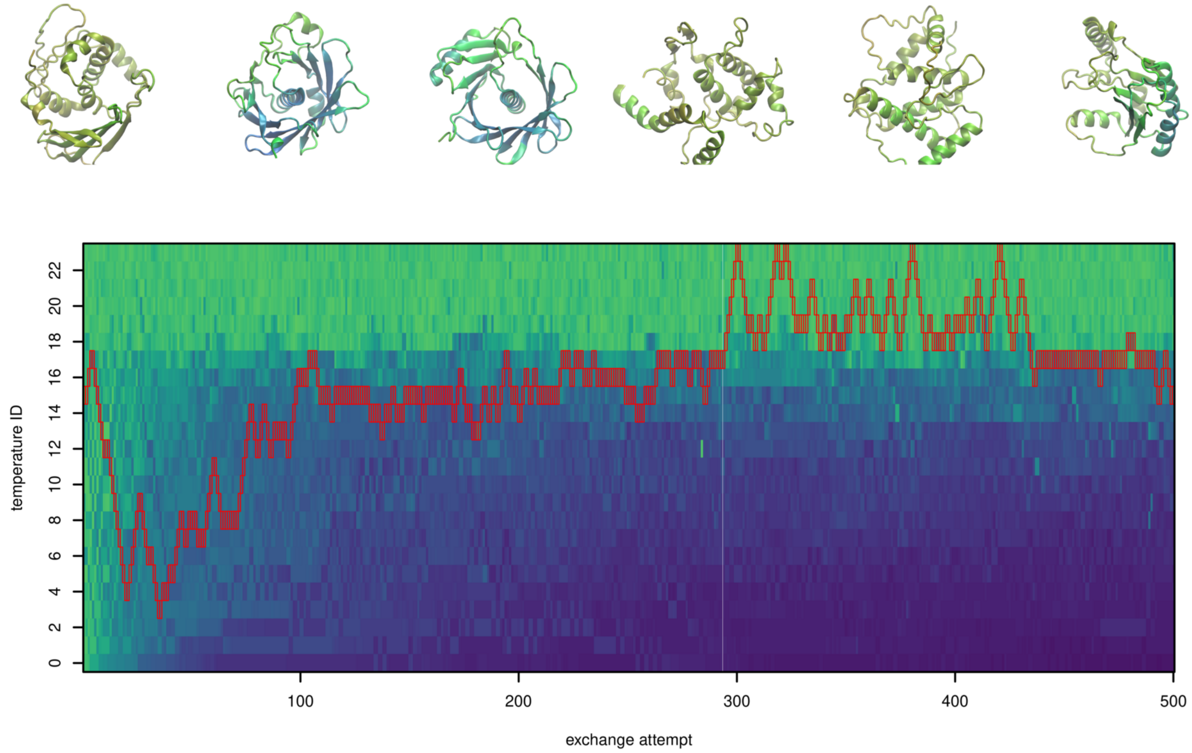


**FIGURE S34** Demultiplexed replica 15 of design of 200-residue proteins by parallel tempering.

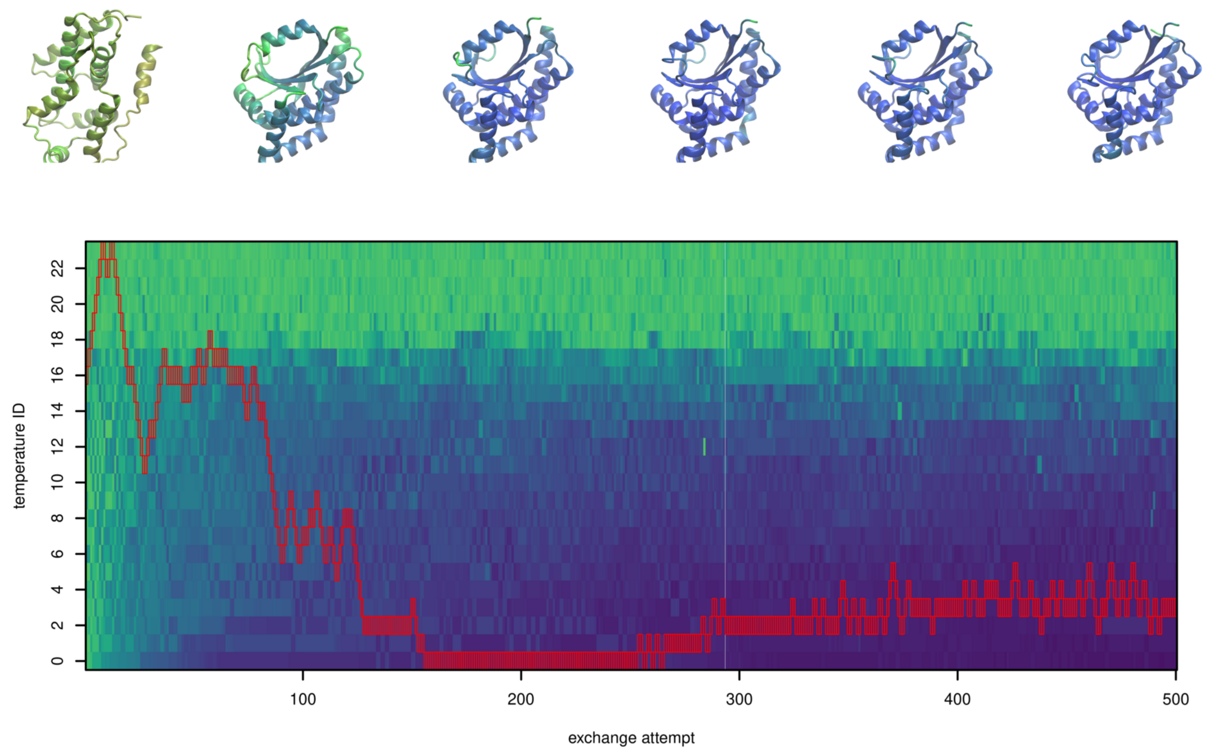


**F I G U R E S35** Demultiplexed replica 16 of design of 200-residue proteins by parallel tempering.

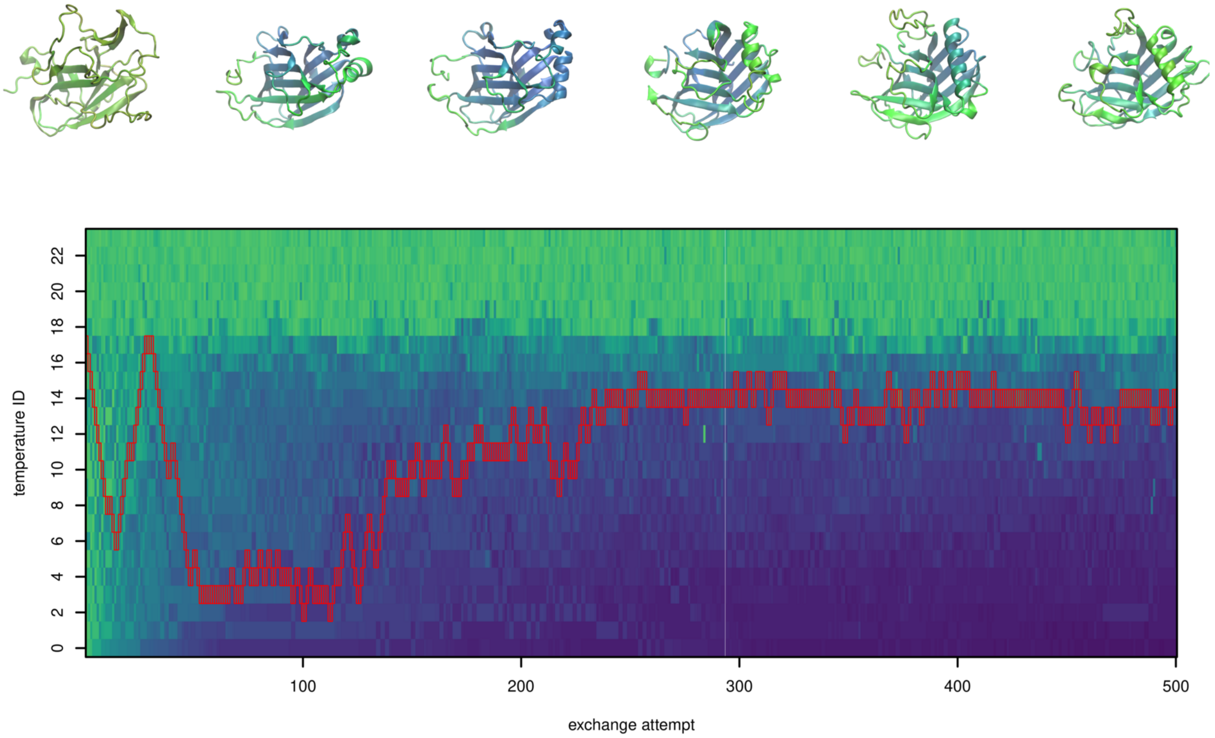


**F I G U R E S36** Demultiplexed replica 17 of design of 200-residue proteins by parallel tempering.

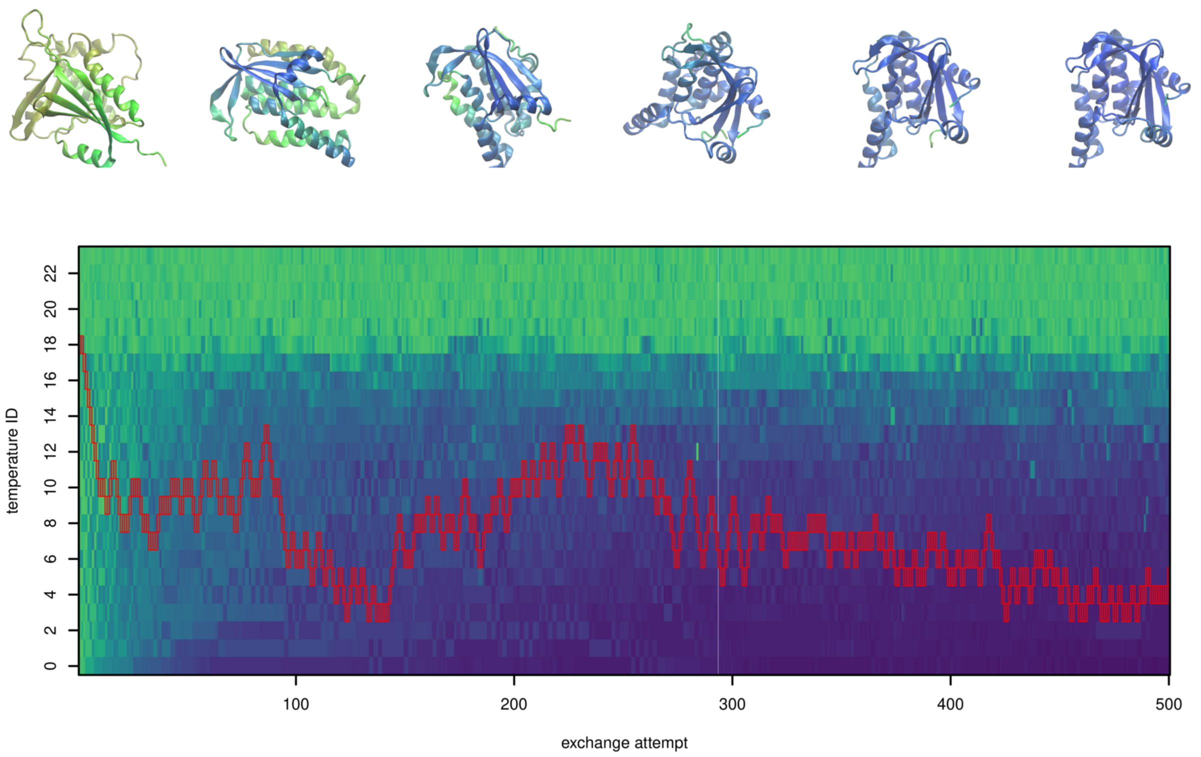


**FIGURE S37** Demultiplexed replica 18 of design of 200-residue proteins by parallel tempering.

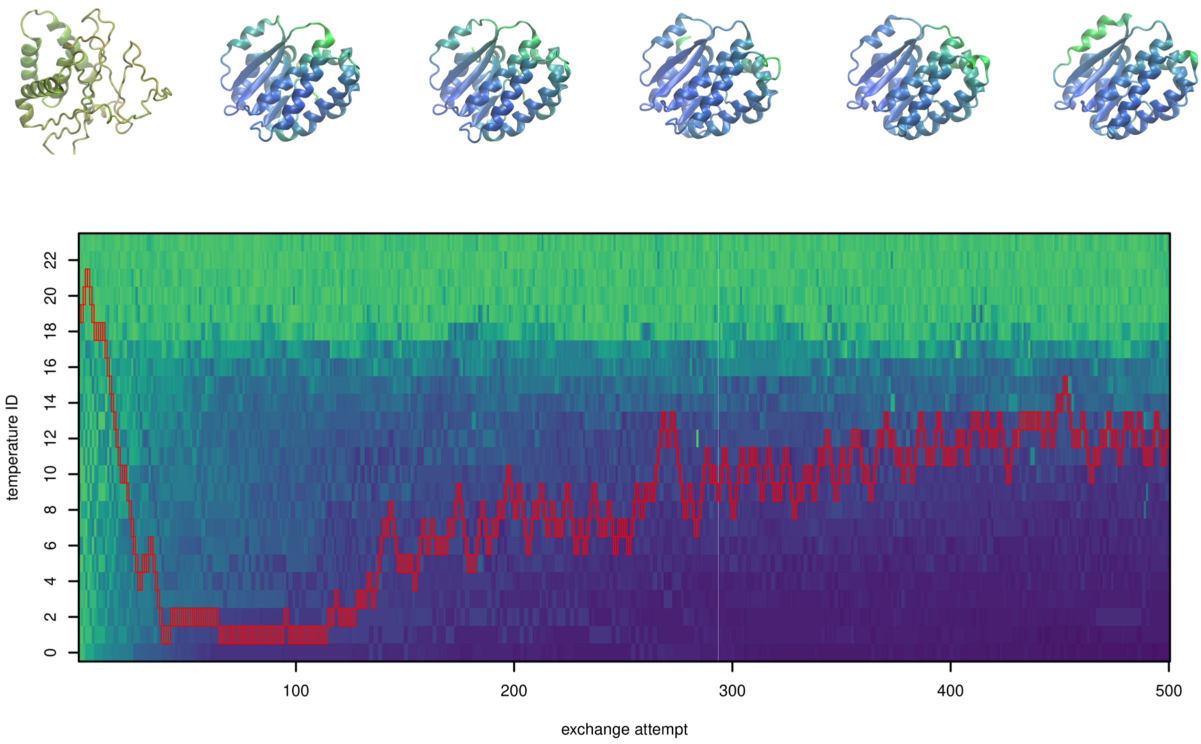


**FIGURE S38** Demultiplexed replica 19 of design of 200-residue proteins by parallel tempering.

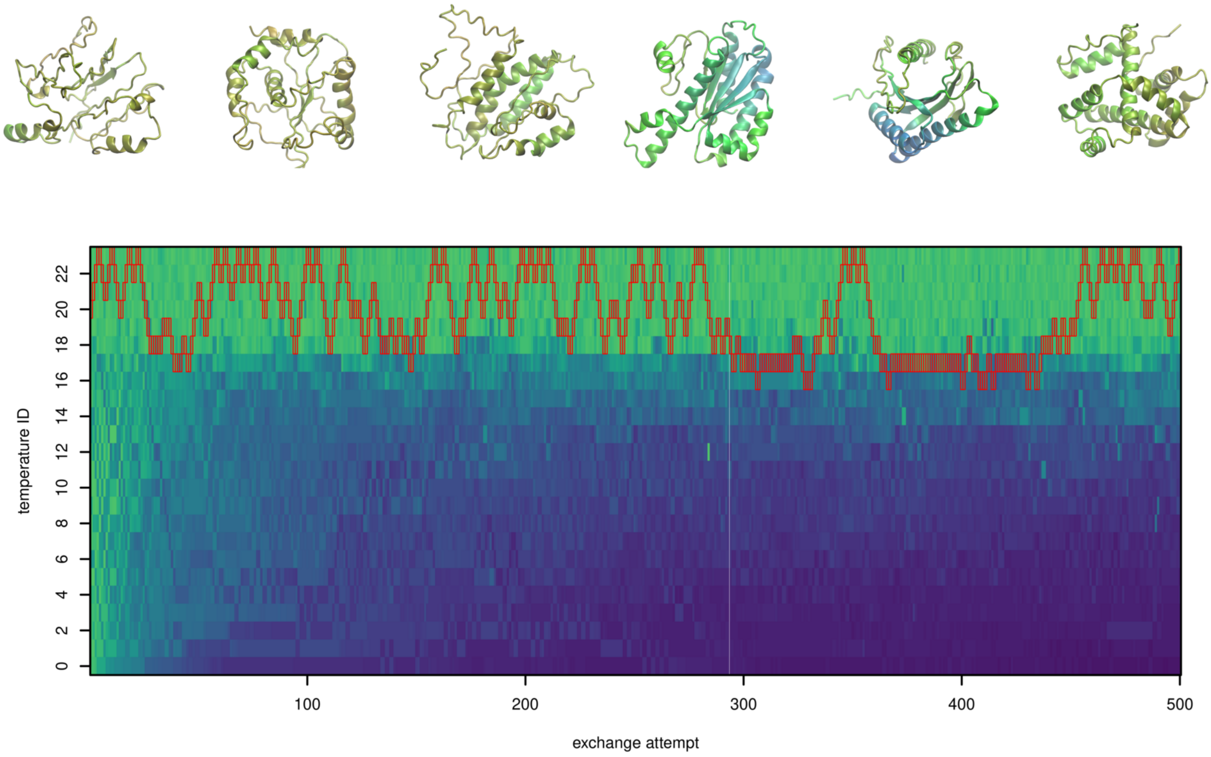


**FIGURE S39** Demultiplexed replica 20 of design of 200-residue proteins by parallel tempering.

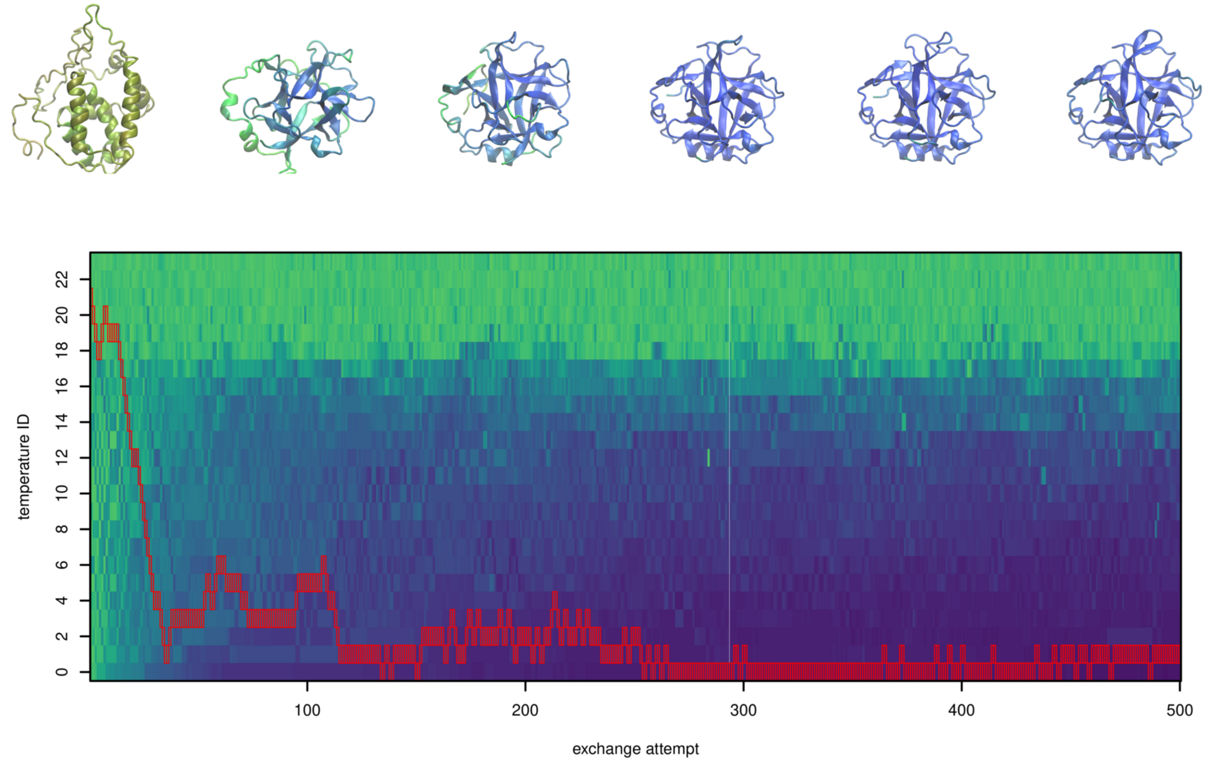


**FIGURE S40** Demultiplexed replica 21 of design of 200-residue proteins by parallel tempering.

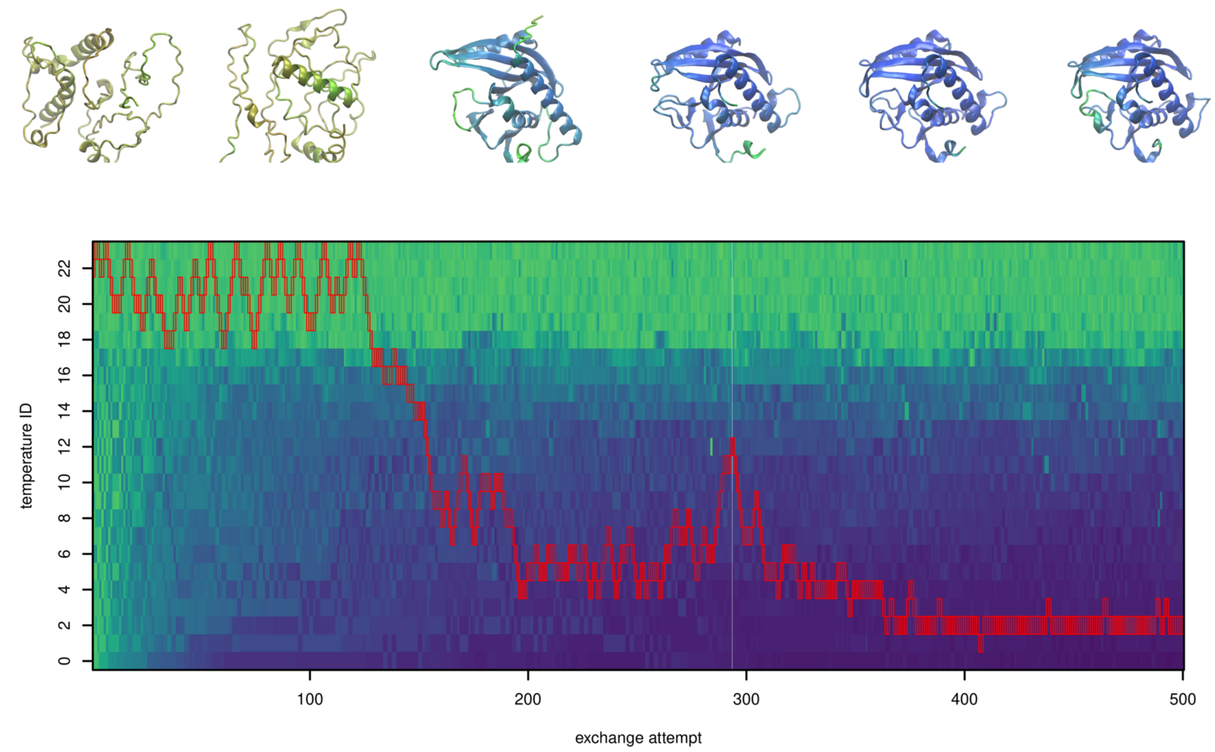


**FIGURE S41** Demultiplexed replica 22 of design of 200-residue proteins by parallel tempering.

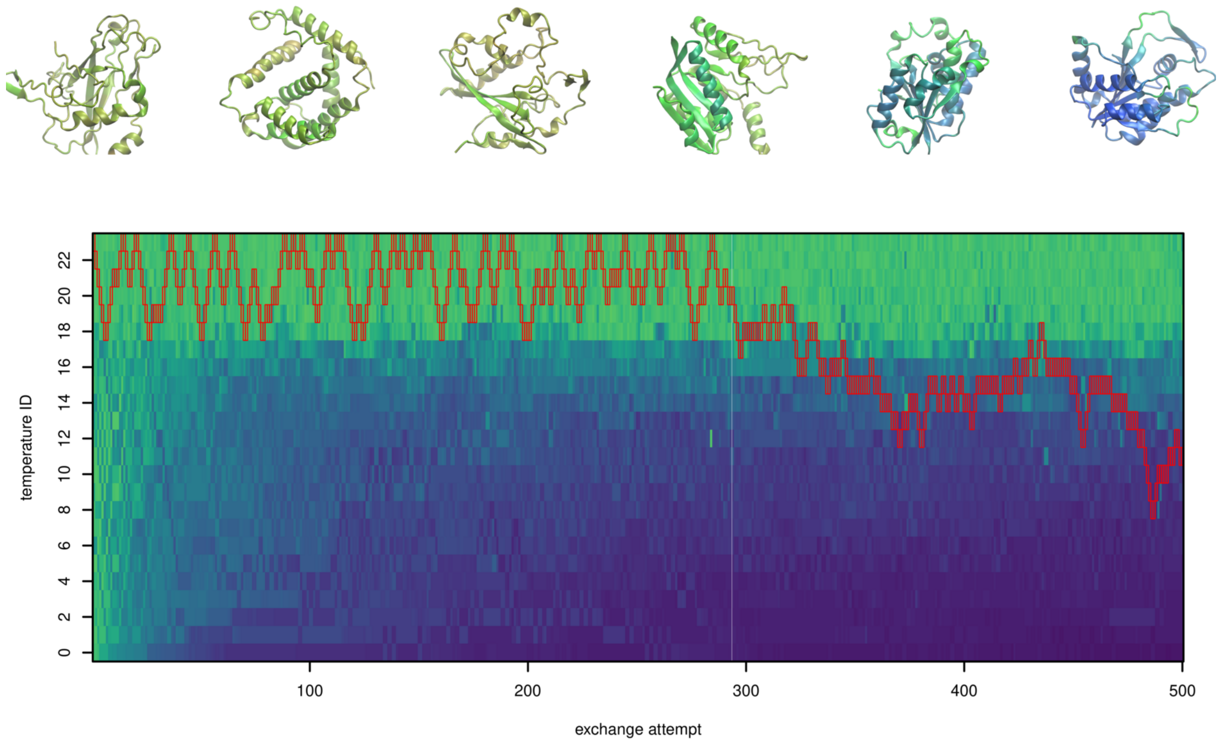


**FIGURE S42** Demultiplexed replica 23 of design of 200-residue proteins by parallel tempering.
